# Distinct dynamics and proximity networks of hub proteins at the prey-invading cell pole in a predatory bacterium

**DOI:** 10.1101/2023.11.29.569176

**Authors:** Ophélie Remy, Yoann Santin, Veronique Jonckheere, Coralie Tesseur, Jovana Kaljević, Petra Van Damme, Géraldine Laloux

## Abstract

In bacteria, cell poles function as subcellular compartments where proteins localize during specific lifecycle stages, orchestrated by polar “hub” proteins. Whereas most described bacteria inherit an “old” pole from the mother cell and a “new” pole from cell division, polarizing cells at birth, non-binary division poses challenges for establishing cell polarity, particularly for daughter cells inheriting only new poles. We investigated polarity dynamics in the obligate predatory bacterium *Bdellovibrio bacteriovorus*, proliferating through filamentous growth followed by non-binary division within prey bacteria. Monitoring the subcellular localization of two proteins known as polar hubs in other species, RomR and DivIVA, revealed RomR as an early polarity marker in *B. bacteriovorus*. RomR already marks the future anterior poles of the progeny during the predator’s growth phase, in a define time window closely following the onset of divisome assembly and the end of chromosome segregation. In contrast to RomR’s stable unipolar localization in the progeny, DivIVA exhibits a dynamic pole-to-pole localization. This behaviour changes shortly before division of the elongated predator cell, where DivIVA accumulates at all septa and both poles. *In vivo* protein interaction networks for DivIVA and RomR, mapped through endogenous miniTurbo-based proximity labeling, further underscore their distinct roles in cell polarization and the importance of the anterior “invasive” cell pole in prey-predator interactions. Our work emphasizes the strict spatiotemporal coordination of cellular processes underlying *B. bacteriovorus* proliferation, offering insights into the subcellular organization of bacteria with filamentous growth and non-binary division.

## Introduction

The cellular content of bacteria is highly organized despite the usual absence of intracytoplasmic organelles. The cell poles constitute particular regions in non-spherical bacteria where a diversity of proteins specifically localize during part or whole of the cell cycle. To polarize diverse functions, bacteria rely on hub proteins that act as scaffolds for the recruitment of other proteins. These polar organizers recognize the cell poles thanks to various features specific to these regions (reviewed in (1)). While cell division typically generates asymmetry in the daughter cells inheriting one “old” pole from the mother cell and one “new” pole from the division site, this does not apply to bacteria producing more than two daughter cells by non-binary division (2). Here, a fraction of the progeny inevitably receives two “new” poles, implying that other mechanisms polarize the offspring. Whereas the organization of the cell poles and the function of polar hub proteins has been the focus of many studies, these investigations mostly used binary-dividing species as model organisms.

*Bdellovibrio bacteriovorus* is a delta-proteobacterium that grows by filamentation and divides in a non-binary manner inside another diderm bacterium, hence featuring an obligate predatory lifestyle. Its lifecycle is commonly divided into two main stages (3, 4). In the attack phase (AP), predators are highly polarized cells (5). A unipolar flagellum propels a fast swim in the milieu. Opposite the flagellated pole, pili are extruded at the so-called “invasive pole” to facilitate attachment to a prey followed by the incursion of the predator cell through the prey outer membrane and peptidoglycan (6, 7). The chromosome of *B. bacteriovorus* is also polarized, featuring a longitudinal *ori-ter* organization with *ori* strictly localized near the invasive pole. Once inside the prey (called bdelloplast), the predatory bacterium starts to elongate as it digests prey cellular contents. During this growth phase (GP), *B. bacteriovorus* replicates its chromosome asynchronously, resulting in odd or even copy numbers corresponding to the number of daughter cells (8). The duration of the GP, the timing of non-binary cell division, and therefore, the number of predator offspring, are determined by prey cell size and composition (9). When exiting the prey, newborn predators display the same polarity as before prey invasion (5, 8). How and when different cellular structures are assigned to the right pole in the numerous future daughter cells remains mysterious.

*B. bacteriovorus* encodes homologs of two proteins described as polar hubs in other species, RomR and DivIVA. DivIVA uniquely localizes at cell poles and septa through its propensity to self-assemble at negatively curved membranes (10, 11), where it plays pivotal roles in polarizing and coordinating various biological events depending on the species, including cell division, chromosome segregation, and sporulation (reviewed in (12)). Mostly present and characterized in monoderm species, DivIVA was recently reported in *B. bacteriovorus* (13) but its precise role, particularly in polarity, remains to be elucidated.

RomR is a key component of the polarity module underlying the directional motility crucial for the predatory behaviour of *Myxococcus xanthus* (14). In this bacterium, RomR (together with RomX (15)) recruits MglA and maintains the active GTP-bound state of this motility regulator at the anterior pole. There, MglA recruits SgmX to stimulate pili extension and retraction (16, 17), pulling the cell forward. The posterior pole predominantly hosts MglB, which converts MglA (with the help of RomY (18)) to the inactive GDP-bound form (19). Intriguingly, RomR-SgmX and MglA-SgmX interactions were reported for the *B. bacteriovorus* homologs (20). Although their role still needs to be clarified in *B. bacteriovorus*, these proteins are required for prey invasion, and RomR is essential even in a mutant strain that grows independently of prey, highlighting its importance for the *B. bacteriovorus* lifecycle (20). RomR also interacts with CdgA (20), a potential invasive-pole receptor for the secondary messenger cyclic-di-guanosine monophosphate (c-di-GMP), proposed to be implicated in various predatory processes in *B. bacteriovorus* (20–22). The localization of RomR at the invasive cell pole of *B. bacteriovorus* during the attack phase (8, 20, 23) supports a role in polarizing predation-related functions. However, the precise subcellular dynamics of RomR during the predatory lifecycle and the extent of its interaction network are still unknown.

Here we set out to determine the subcellular localization of RomR and DivIVA throughout the *B. bacteriovorus* cell cycle, leveraging single-cell reporters of key cellular processes imaged by live-cell fluorescence microscopy. The distinct localization patterns of these proteins reveal a precise order of events during the predatory growth phase. Our study sheds light on the establishment of cell polarity by highlighting RomR as an early marker of the future invasive poles prior non-binary cell division in *B. bacteriovorus*. Using a proximity labelling approach for the first time in this bacterium, we connect DivIVA and RomR to specific protein networks, refining our understanding of the role of polarity in bacterial predation.

## Results

### RomR marks the invasive cell poles of future B. bacteriovorus progenies during the growth phase

To gain spatiotemporal insights into the localization of RomR during the predatory lifecycle of *Bdellovibrio bacteriovorus*, we constructed a strain expressing a fusion of RomR to mCherry as single copy from its native chromosomal locus (as in (24)), in an otherwise wild-type background (*romR*::*romR-mcherry*). We imaged these cells by phase contrast and fluorescence microscopy, during the attack phase and upon infection of *E. coli* (**Fig 1A, Supp Fig 1A**). Consistent with previous reports using RomR fluorescent fusions expressed from a replicative (8) or chromosomally integrated plasmid (20, 23), or from the native locus (24), attack-phase *B. bacteriovorus* cells featured a single RomR-mCherry focus at the invasive cell pole (8, 20). *B. bacteriovorus* cells retained their single RomR-mCherry focus upon prey invasion (**Fig 1A, Supp Fig 1A**). Time-lapse imaging of these bdelloplasts showed that during the early growth phase, a second focus appeared at the pole opposite the first one, in about half of the cases (**Fig 1A**, dark pink arrowhead, **Supp Fig 1A-B**). In all bdelloplasts, additional RomR-mCherry foci formed later within the *B. bacteriovorus* elongating cell body (**Fig 1A**, light pink arrowheads, **Supp Fig 1A**). Notably, these new RomR-mCherry foci did not move away from their initial subcellular position, indicating that they directly accumulate at their final destination. Moreover, all RomR foci marked (future) constriction sites (as seen by phase contrast, asterisks in **Fig 1A**, **Supp Fig 1A**), but not all septa were labelled with a RomR focus at the time of division.

**FIG 1.**
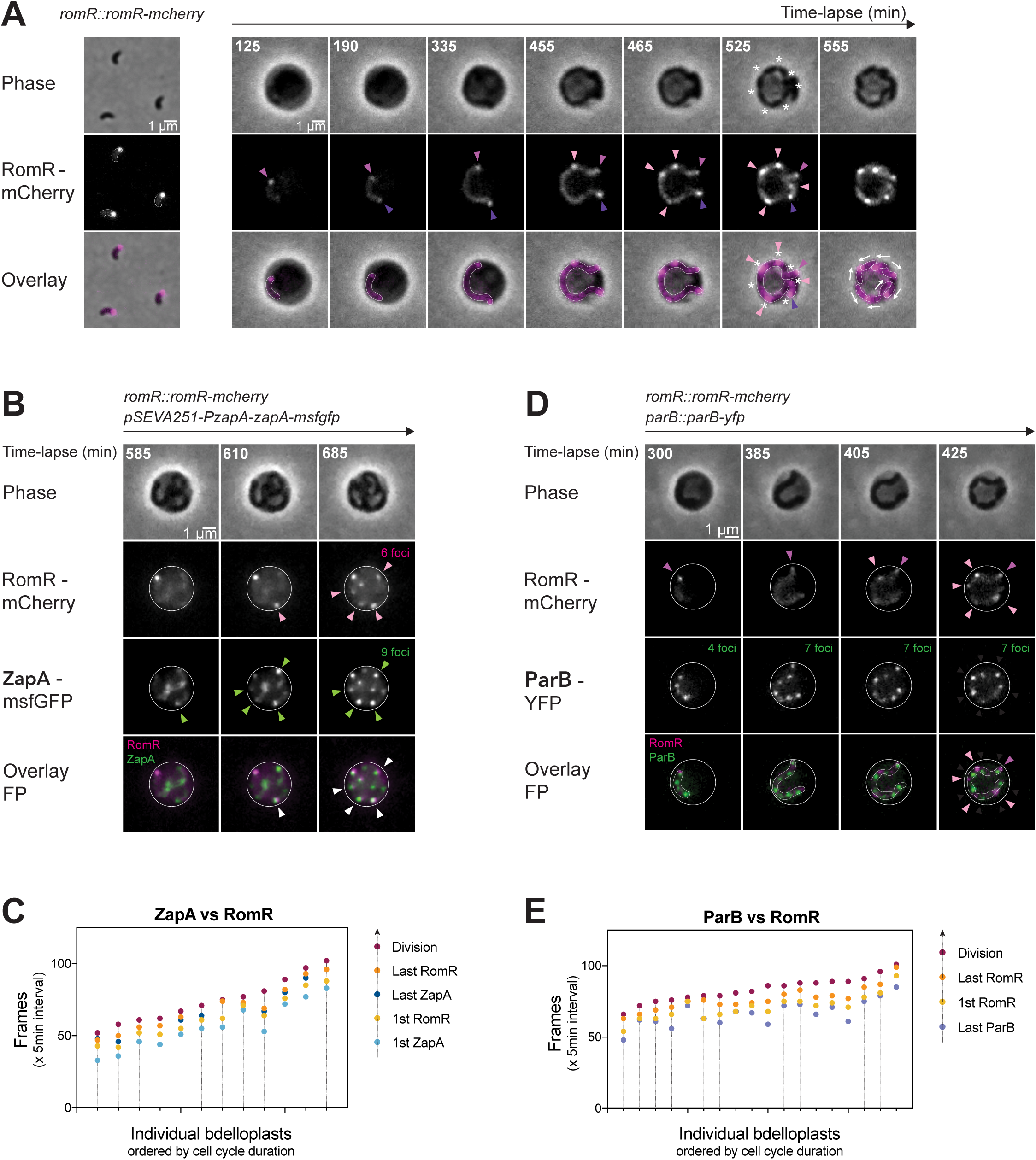
The appearance of new RomR foci follows a precise spatiotemporal pattern during the *B. bacteriovorus* growth phase. **(A)** Spatial localisation of RomR-mCherry in attack (AP) and growth phase (GP) cells. Left: Representative phase contrast and fluorescence images of AP cells for the *B. bacteriovorus* HD100 *romR::romR-mcherry* strain (GL1466). Right: Representative bdelloplast containing a GP cell of the *romR::romR-mcherry* strain (GL1466). Predators were mixed with exponentially grown MG1655 *E.coli* prey for 90 min prior imaging in time-lapse with 5 min intervals. Both: From top to bottom, phase contrast channel, mCherry channel, and overlay of both channels. Dark pink arrowheads point to the first RomR-mCherry focus; purple arrowheads to the second focus seen during growth; light pink arrowheads to the new foci at the end of the growth phase; asterisks to septa; arrows to the RomR foci orientation relative to *B. bacteriovorus* daughter cells. **(B-C)** Temporal localisation of RomR-mCherry compared to the early divisome component ZapA-msfGFP (GL2378). (B) Representative bdelloplast containing a GP cell of *B. bacteriovorus* GL2378 strain mixed with exponentially grown *E. coli* GL522 prey (carrying the *mreB_L251R_* mutation and therefore larger than the wild-type MG1655 strain) for 90 min prior imaging, with 5 min intervals. From top to bottom, phase contrast channel, mCherry channel, GFP channel and overlay of both FP channels with RomR-mCherry in magenta and ZapA-msfGFP in green. Pink arrowheads point to new RomR-mCherry foci; green arrowheads to new ZapA-msfGFP foci; white arrowheads to the colocalized foci. The final number of foci of RomR and ZapA are written in magenta and green respectively. (C) Graphical representation of the sequential timing of foci appearance in GP. For each bdelloplast, 5 timepoints (frames) were manually recorded and plotted according to the color code shown on the right: (1) when the first ZapA-msfGFP foci appeared, (2) when the last ZapA-msfGFP foci appeared, (3) when new RomR-mCherry foci are seen (in addition to the focus already present during attack phase and prey invasion), (4) when the last RomR foci appeared, and (5) when daughter cells physically separated (division of the mother cell). Individual bdelloplasts are distributed along the X axis, in ascending order of cell cycle duration. **(D-E)** Temporal localisation of RomR-mCherry compared to the chromosome segregation protein ParB-YFP (GL1655). (D) Representative bdelloplast containing a GP cell of *B. bacteriovorus* GL1655 strain mixed with exponentially grown *E. coli* GL522 prey for 90 min prior imaging, with 5 min time intervals. From top to bottom, phase contrast channel, mCherry channel, YFP channel and overlay of both FP channels with RomR-mCherry in magenta and ParB-YFP in green. Dark pink arrowheads point to the old RomR-mCherry focus; light pink to the new RomR-mCherry foci. The number of ParB foci is written in green. (E) Graphical representation of the sequential timing of foci appearance in GP. For each bdelloplast, 4 timepoints (frames) were manually recorded and plotted according to the color code shown on the right: (1) when the last ParB-YFP focus appeared, (2) when new RomR-mCherry foci are seen, (3) when the last RomR foci appeared, and (4) when daughter cells physically separated (division of the mother cell). Individual bdelloplasts are distributed along the X axis, in ascending order of cell cycle duration. *B. bacteriovorus* filament and bdelloplasts outlines were drawn manually based on the fluorescence and phase contrast images.

Whereas the positioning of RomR foci did not seem to follow a predictable pattern among the observed bdelloplasts, we noticed common features in their localization dynamics (**Fig 1A**). First, new RomR-mCherry foci formed when the *B. bacteriovorus* cell was still elongating and before all constriction sites were visible. Second, no additional RomR-mCherry focus appeared after the physical separation of daughter cells. Observations of bdelloplasts right after division showed that each daughter cell inherited precisely one polar RomR-mCherry focus from the corresponding constriction site in the filamentous mother cell. Consistently, we never observed two consecutive septa without a RomR focus. These findings imply that the positioning of RomR during *B. bacteriovorus* growth is highly regulated to result in one focus per cell. Since RomR marks the invasive cell pole of attack phase predators and directly localizes at the future invasive pole of daughter cells along the growing predator cell, our data collectively suggest that RomR is an early cell polarity marker in *B. bacteriovorus*.

### RomR foci appear within a specific time window during the B. bacteriovorus late growth phase

Since all new RomR-mCherry foci appeared before the division event in *B. bacteriovorus*, we sought to shed light on the timing of RomR accumulation relative to the predator’s cell cycle by monitoring the assembly of the division machinery and chromosome replication and segregation. ZapA is a highly conserved protein known to stabilize the early assembly of FtsZ protofilaments (25). Its accumulation at future division sites was used as a marker for early divisome assembly in several species (26–29). Time-lapse imaging of a ZapA-msfGFP fusion (expressed from the endogenous *zapA* promoter on a replicative plasmid) in the *romR::romR-mcherry* strain revealed that both proteins form foci that appear during the late growth phase in *B. bacteriovorus* (**Fig 1B**). However, ZapA foci always appeared first, followed by RomR foci (**Fig 1B-C**, green and pink arrowheads, respectively). Consistent with our previous observations, the last RomR focus appeared before the physical separation of daughter cells (**Fig 1B-C**). Moreover, each new RomR-mCherry focus colocalized with a ZapA-msfGFP focus (**Fig 1B**, white arrowheads), but the growing predator cell contained fewer RomR foci than ZapA foci (**Fig 1B**). This confirms that RomR accumulates at some, but not all, division sites in *B. bacteriovorus*. Similar co-localization patterns were obtained when a second conserved early divisome component, FtsA, which anchors FtsZ in many bacterial species (30), was labelled together with RomR (**Supp Fig 1C**). Next, we tracked the progress of chromosome replication and segregation by monitoring ParB foci, which label the origin of replication of each copy of the chromosome during the *B. bacteriovorus* growth phase (8), together with RomR-mCherry. We found that new RomR foci only appeared after the detection of the last ParB focus, i.e., after the initiation of the last round of chromosome replication and segregation (**Fig 1D-E**). However, the first new RomR-mCherry spots were seen shortly after the appearance of the last ParB focus. Altogether, our results highlight that RomR marks future invasive cell poles by localizing at a subset of future constriction sites after the onset of divisome assembly and during a precise time window of the *B. bacteriovorus* growth phase, i.e., at the transition between the end of chromosome segregation and the release of the predatory progeny.

### DivIVA preferentially localises at the invasive cell pole during the attack phase

To extend our exploration of potential polarity markers in *B. bacteriovorus*, we investigated the localization dynamics of DivIVA. Snapshots of attack phase cells natively expressing *divIVA-msfgfp* from the chromosomal locus showed that DivIVA-msfGFP displayed a polar localization (**Fig 2A**), consistent with previous reports (13). However, DivIVA-msfGFP was not evenly distributed between the two cell poles in *B. bacteriovorus*, in contrast with characterized DivIVA homologues (10, 11, 31, 32). Instead, DivIVA was unipolar in about one-third of the attack phase *B. bacteriovorus* cells (38.15%, n = 4314), whereas the majority of cells contained two DivIVA foci of distinct fluorescence intensities (**Fig 2A-B**). Using FM4-64, which stains the membrane-sheathed flagellum of *B. bacteriovorus* (8), we noticed that DivIVA preferentially accumulates at the invasive cell pole (**Fig 2B**). Quantified comparisons of DivIVA-msfGFP unipolar localization patterns revealed that DivIVA localises almost twice as much at the invasive pole than at the flagellated pole (**Fig 2B**). Demographs of the DivIVA-msfGFP fluorescence signal confirmed this trend when cells were oriented based on other markers of the invasive pole, either RomR (**Fig 2C**) or *ori* (8) (**Supp Fig 2A**). When expressed in *E. coli,* DivIVA-msfGFP localized similarly as reported for previously characterized DivIVA homologs, clearly delineating both poles and septa in predivisional cells (**Supp Fig 2B**). These results indicate that, besides its spontaneous accumulation at curved membranes, DivIVA is subjected to a *B. bacteriovorus*-specific regulation of its localization in attack phase cells. To further assess DivIVA-msfGFP dynamics in *B. bacteriovorus,* we performed short one-minute interval time-lapses on attack phase cells. Compared to RomR which remained stable at one pole throughout the experiment, DivIVA foci appeared to alternate between cell poles and unipolar vs. bipolar states (**Fig 2D**). This dynamic pattern was also observed during the transition phase between the invasion of the prey’s periplasm and the initiation of *B. bacteriovorus* cell elongation (**Fig 3A**, see time points 65 to 127.5 min). In summary, DivIVA switches from pole to pole during the non-proliferative phase of the *B. bacteriovorus* cell cycle, with a bias towards the invasive cell pole.

**FIG 2.**
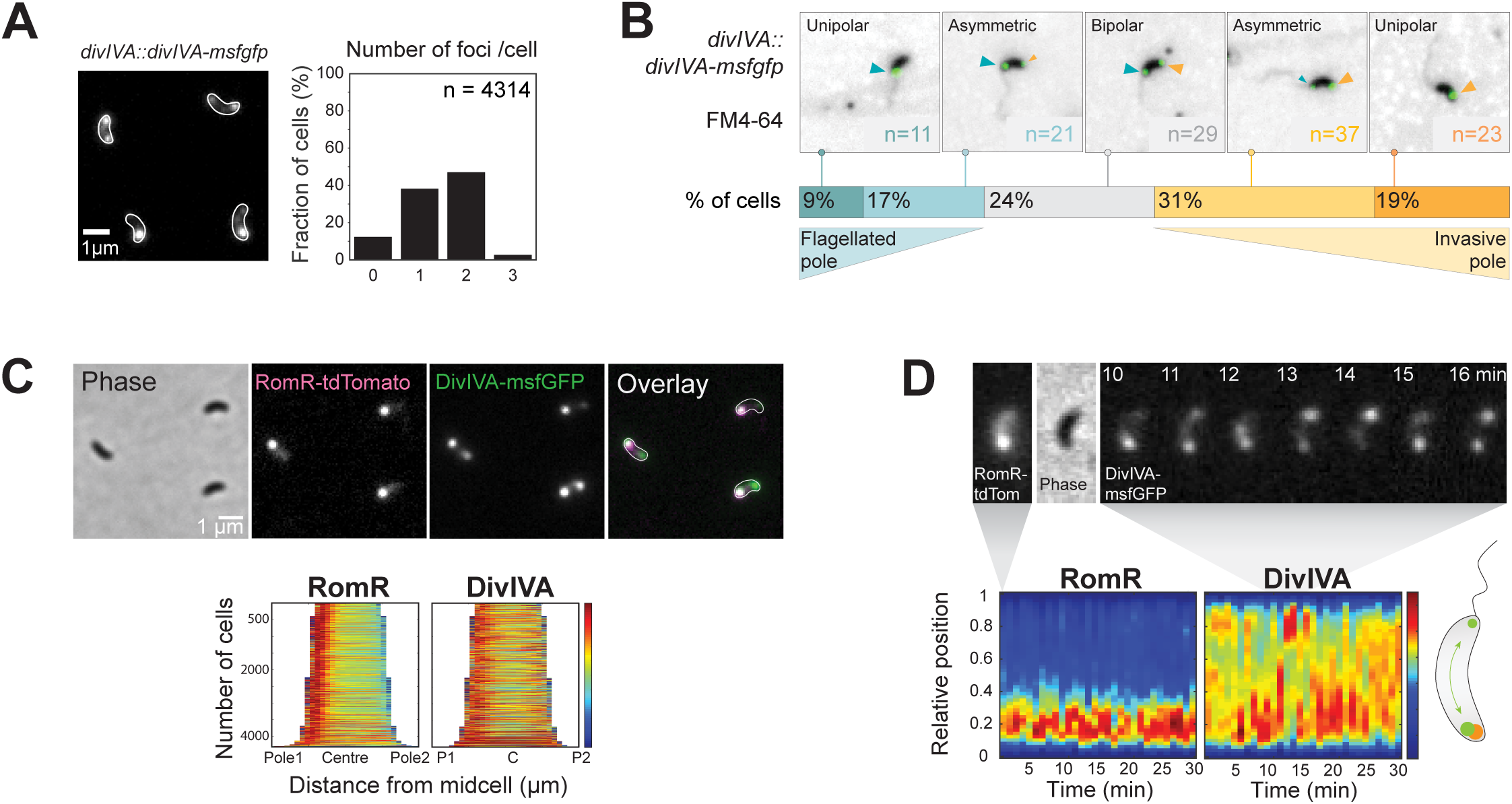
DivIVA polar localisation is dynamic during the attack phase. **(A)** DivIVA-msfGFP does not always uniformly localize at both poles in *B. bacteriovorus*. Left: Representative GFP channel image of attack phase *B. bacteriovorus divIVA*::*divIVA-msfgfp* (strain GL1620) with cell outlines obtained with Oufti based on the corresponding phase contrast image. Right: Histogram representing the percentage of *B. bacteriovorus* AP cells with 0, 1, 2 or 3 foci of DivIVA-msfGFP; n indicates the number of cells analysed in a representative experiment. **(B-C)** DivIVA-msfGFP localizes preferentially at the non-flagellated pole, identified using the FM4-64 membrane dye (B) or RomR-Tdtomato (C). (**B**) Top: Representative images of *divIVA::divIVA-msfgfp* AP cells (GL1620) stained with FM4-64, labelling the outer membrane and the sheathed flagellum of *B. bacteriovorus*. n indicates the number of cells analysed in a representative experiment. Bottom: Schematics representing the percentage of each indicated category of DivIVA-msfGFP localization patterns: (1) Unipolar at the flagellated pole, (2) Asymmetric bipolar at the flagellated pole, (3) Symmetric bipolar, (4) Asymmetric bipolar at the invasive pole, and (5) Unipolar at the invasive pole. (**C**) Top: Representative images of *divIVA::divIVA-msfgfp* / *pTNV215-romR-tdtomato* AP cells (GL944). From left to right: Phase, mCherry, GFP channels and overlay of the fluorescent channels with RomR-tdTomato in magenta and DivIVA-msfGFP in green. Bottom: Demographs of the fluorescent signals in a population of cells of the same strain sorted by length and oriented with the highest RomR-tdTomato signal intensity on the left, labelled as pole 1. The blue-to-red heatmap indicates low-to-high intensities. **(D)** Representative AP cell of the same strain as in C during a short time-lapse with 1 min time intervals. Top: left, mCherry channel at time 0 min, middle is the phase contrast channel at time 0 min; right, GFP channel from 10 to 16 min. Bottom: kymographs of the same cell with the relative fluorescence intensities of RomR-tdTomato (left) and DivIVA-msfGFP (right); the blue-to-red heatmap indicates low-to-high intensities; for both kymographs, the cell is oriented with the highest RomR-tdTomato intensity at the bottom. A schematic representation (oriented similarly) of the foci position and dynamics is shown on the bottom right.

**FIG 3.**
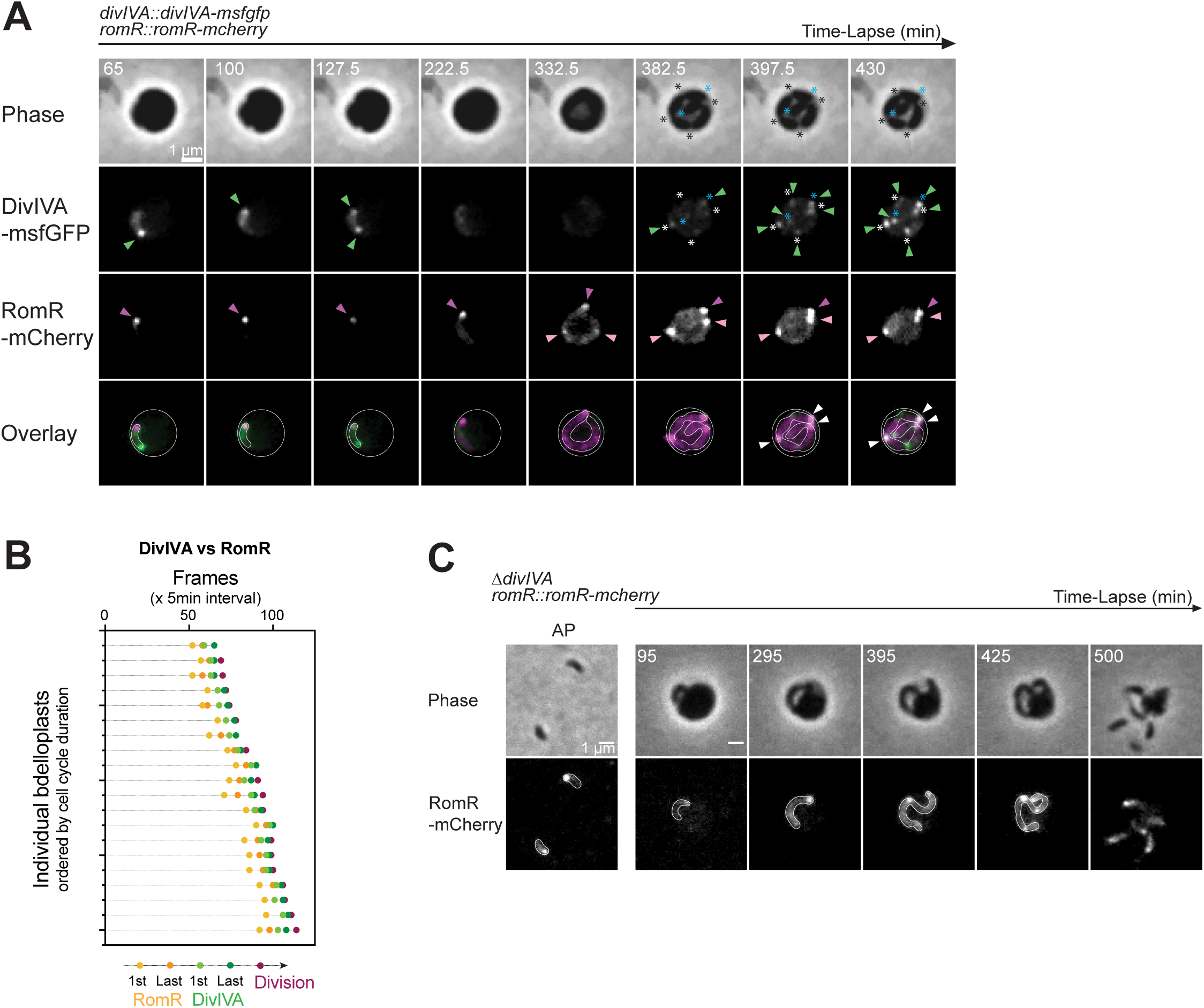
RomR and DivIVA localise differently during the growth phase. **(A-B)** RomR-mCherry new foci appear earlier than DivIVA-msfGFP at the end of the growth phase of *B. bacteriovorus*. (**A**) Representative bdelloplast containing a GP *B. bacteriovorus* cell (strain GL1471). Predators were mixed with exponentially grown GL655 *E. coli* prey for 45 min prior time-lapse imaging with 2.5 min time intervals. From top to bottom: Phase contrast, GFP, mCherry channels and overlay of both FP channels with RomR-mCherry in magenta and DivIVA-msfGFP in green. Dark pink arrowheads point to the old RomR-mCherry focus; light pink arrowheads to the new ones; green arrowheads to DivIVA-msfGFP foci; white arrowheads to the colocalized foci. White and black asterisks indicate the visible constriction sites and blue asterisks the two poles of the mother cell. (**B**) Graphical representation of the sequential timing of foci appearance in GP for this strain. For each bdelloplast, 5 timepoints (frames) were manually recorded and plotted according to the color code indicated on the right: (1) appearance of new RomR-mCherry foci, (2) appearance of last RomR-mCherry foci, (3) new DivIVA-msfGFP foci, (4) last DivIVA-msfGFP foci, and (5) the physical separation of daughter cells (division of the mother cell). Individual bdelloplasts are distributed along the Y axis, in ascending order of cell cycle duration. **(C)** RomR-mCherry localization is not modified in a Δ*divIVA* strain in attack (AP) and growth phase (GP) cells. Left: Representative phase contrast and fluorescence images of AP cells for the Δ*divIVA romR::romR-mcherry* strain (GL2380). Right: Representative bdelloplast containing a GP cell of the same strain. *B. bacteriovorus* were mixed with exponentially grown MG1655 *E.coli* prey for 90 min prior imaging in time-lapse with 5 min intervals. Each time-lapse experiment was performed at least 3 times. *B. bacteriovorus* cells and bdelloplasts outlines were drawn manually based on the fluorescence and phase contrast images.

### DivIVA and RomR display distinct subcellular localization dynamics during B. bacteriovorus growth

Considering the preferential accumulation of DivIVA-msfGFP at the invasive pole during the attack phase, we followed its localisation throughout the *B. bacteriovorus* growth phase within the prey in comparison with RomR-mCherry, in a strain expressing both fluorescent fusions as single copies from their native loci (**Fig 3A**). During predator elongation, the DivIVA-msfGFP signal disappeared, whereas the RomR-mCherry focus remained clearly visible (**Fig 3A, Movie S1**). DivIVA was still undetectable when additional RomR foci appeared. As shown by immunoblotting (**Supp Fig 2C**), the limited amount of DivIVA proteins during growth likely prevents the nucleation of DivIVA foci in the elongated predator cell. The disappearance of DivIVA foci during the growth phase was not observed in a previous study (13), possibly because smaller prey was employed, resulting in a reduced predator growth phase (9), and the imaging of different bdelloplasts at each timepoint separated by longer time intervals (reproduced in **Supp Fig 2D**). However, in both previously used and our conditions, new DivIVA-msfGFP foci gradually emerged at visible constriction sites (**Fig 3A**, white asterisks) and at cell poles (**Fig 3A**, blue asterisks), towards the end of the growth phase (**Fig 3A, Supp Fig 2B, Movie S1**). Notably, DivIVA foci formed only after all RomR foci were established. They eventually marked every septum (some of them colocalizing with RomR; **Fig 3A**, white arrows) and both poles of the mother cell, shortly before or at the time of cell division (**Fig 3A-B, Movie S1**).

### RomR localization and B. bacteriovorus cell morphology are independent of DivIVA

The biased accumulation of DivIVA at the invasive pole might reflect a role in the attack of the prey. To test this possibility, we deleted the *divIVA* gene from the genome. In contrast with RomR, which is essential for *B. bacteriovorus* (20), the absence of DivIVA did not visibly impact the predator cell. Unlike in a previous study (13), in attack phase, the Δ*divIVA* mutant showed no morphological phenotype in length, width, or curvature, compared to the wild-type strain (**Supp Fig 3A-B**). Additionally, the efficiency of prey killing, measured by the rate of lysis of an *E. coli* population upon addition of *B. bacteriovorus*, was not impacted by the *ΔdivIVA* mutation (**Supp Fig 3C**). As expected from our observation of the relative timing of RomR and DivIVA foci formation (**Fig 3A-B**), RomR-mCherry localisation was unperturbed by the lack of DivIVA (**Fig 3C**).

### Proximity labelling uncovers distinct networks for both RomR and DivIVA polar proteins

Distinct subcellular dynamics of DivIVA and RomR throughout the cell cycle of *B. bacteriovorus* suggest that these proteins play different roles. Both proteins have reported interaction partners in other species, which determine their function (12, 19, 33). Thus, to get insight into the protein landscape of DivIVA and RomR in *B. bacteriovorus*, we applied a proximity-dependent biotinylation (34) using miniTurbo (35), a proximity labelling derivative of BioID. Briefly, the miniTurbo enzyme fused to the protein of interest (POI, bait) uses ATP and biotin (supplemented in the medium) as substrates to produce a short-lived and highly reactive biotinoyl-5’-AMP intermediate, which covalently binds to amine group in lysine residues, located within a ∼10 nm radius, thereby labelling proteins in the close vicinity of the bait or “proxisome” (**Fig 4A**). Biotinylated proteins are isolated on streptavidin beads and identified by mass spectrometry (MS) analysis (see Methods). *E. coli* prey and attack phase *B. bacteriovorus* (producing DivIVA-miniTurbo, RomR-miniTurbo, or a control fusion, see Methods) were incubated with biotin prior predation and then during growth for 5 h, allowing labelling of proximal proteins during the entire predatory lifecycle. Distinct biotinylation patterns were obtained in total protein extracts of the three strains, as detected with a streptavidin conjugate (**Supp Fig 4A**), indicating POI-specific biotin labelling. MS and immunoblotting analyses of total proteome extracts upon miniTurbo-mediated proximity labelling in *B. bacteriovorus*, detailed in the Material and Methods section, showed efficient and specific capture of biotinylated proteins (**Supp Fig 4B**), correlation among triplicate samples (**Supp Fig 4C**), and the significant enrichment of POIs in their respective setups (**Fig 4B-C and Supp Fig 4D**), validating the approach. Overall, the unique proxisomes identified are in line with the idea that RomR and DivIVA participate in different networks and therefore, carry out distinct cellular functions. Further strengthening the significance of our approach, the RomR proximal proteins (**Fig 4B**) include MglA, reported as an indirect RomR partner in *B. bacteriovorus* (20), also known to act in the same polarity network as RomR in *M. xanthus* (19, 33, 36). Additionally, all previously described hits are known to localize at the invasive cell pole of *B. bacteriovorus* (labelled (i) in the table, **Fig 4B**) while several of the remaining proteins were proposed to be important for predation according to a TnSeq analysis in another *B. bacteriovorus* strain (37) (labelled # in the table, **Fig 4B**). Besides, c-di-GMP binding proteins (22) (labelled * in the table, **Fig 4B**) were identified (of note, the diguanylate cyclase localized at the invasive pole, DgcB (Bd0742) (21) was also identified, although above our significance threshold (FDR 0.05)). Finally, several proteins of the identified RomR proxisome are functionally related (e.g., type IV filaments components (38)) or encoded in the same genomic context (e.g., Bd3261 and Bd3262). The identification of proximal proteins residing in known protein complexes was also evident for DivIVA (**Fig 4C**). Here, among the 20 significant hits found in the proxisome, five are annotated as associated to the chemosensory system. Further, the DivIVA proxisome is enriched in post-translational regulatory proteins such as Lon2, DnaK, ClpX, HptG or FtsH1. These data imply that DivIVA might be implicated in a broader and more complex network of interactions, which will be the focus of further studies. To summarize, we applied a biotin-based proximity labelling approach for the first time in *B. bacteriovorus*, which brought new insights into the specific protein networks of RomR and DivIVA.

**FIG 4.**
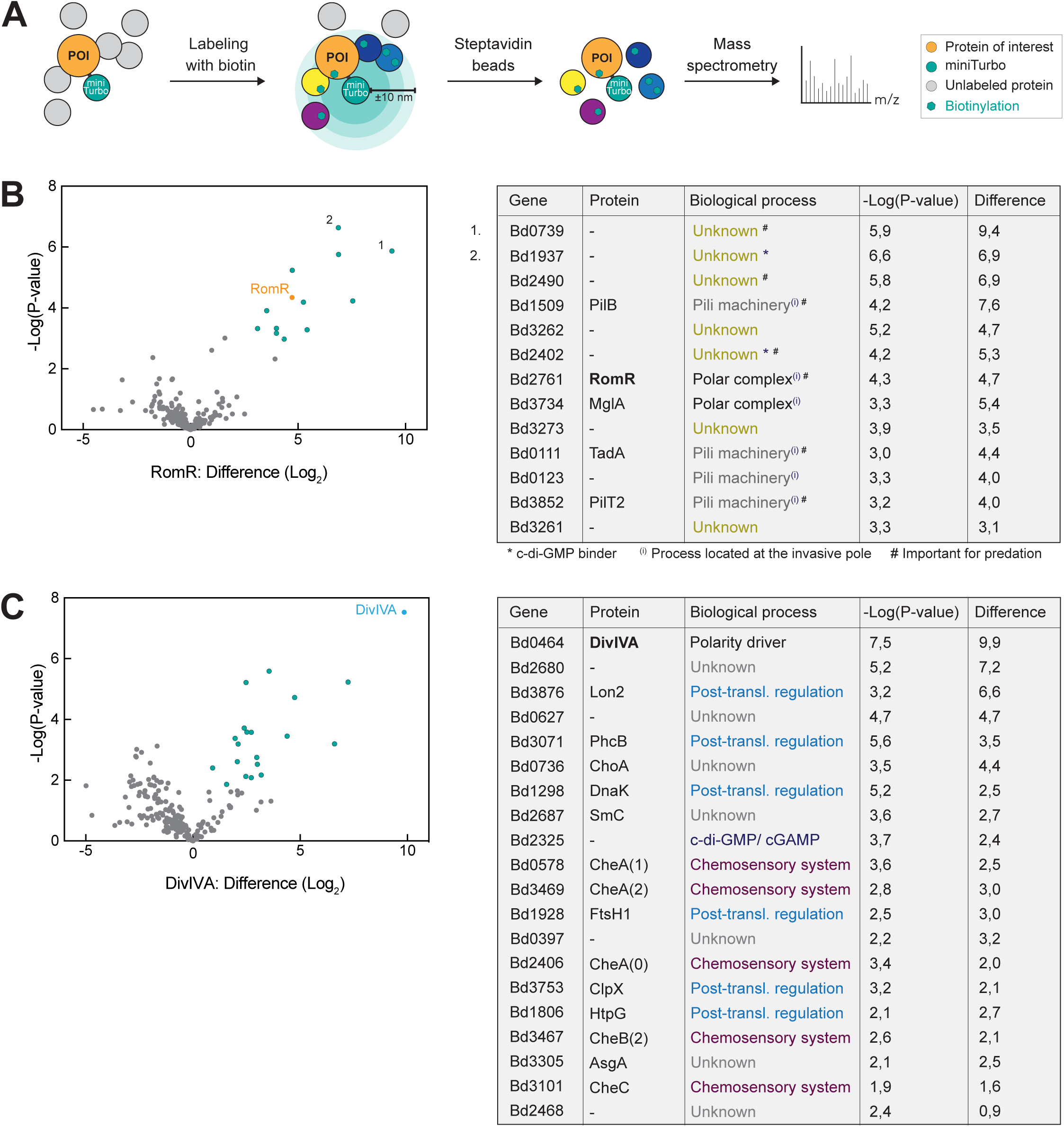
Proximity labelling reveals specific protein networks associated with *B. bacteriovorus* RomR and DivIVA *in vivo*. **(A)** Schematic representation of the proximity labelling assay performed using miniTurbo fused to the protein of interest (POI). Briefly, the POI(-miniTurbo fusion) is expressed from its native locus in the cell. In the presence of biotin, protein labelling occurs (biotinylation). Proteins in the close (∼10 µm) environment (i.e., direct and indirect partners) of the POI are biotinylated, captured using streptavidin beads and identified by MS. **(B-C)** Results of the proximity labelling for RomR-miniTurbo (B) and DivIVA-miniTurbo (C) are plotted in a volcano plot on the left, with the difference (log_2_) of the tested strain compared to the other conditions (i.e., RomR-miniTurbo or DivIVA-miniTurbo, and ParB-miniTurbo) on the X axis and the p-value (-Log(P-value)) on the Y axis. ParB-miniTurbo was used as an additional control condition for the MS identification of POI-specific hits. On the right, the significant hits (gene and protein names; FDR 0.01) are listed, together with the biological process in which they are predicted or known to be implicated.

### RomR directly interacts with a set of essential and uncharacterized proteins

RomR proximity labelling revealed several hits previously implicated in functions related to predatory attack and proposed to occur at the invasive cell pole (e.g., pili, c-di-GMP signalling). Considering the essentiality of RomR, we then characterized this network further. Since proximity labelling also identifies indirect and/or transient partners, we tested direct interactions between RomR and selected proximal proteins, using the PopZ-linked Apical Recruitment (POLAR) assay in *E. coli* (39) (which lacks homologs of the polarity module proteins RomR and MglA). Specifically, RomR is fused to msfGFP and PopZ_H3H4_, the oligomerization domain H3H4 of PopZ, a *Caulobacter crescentus* protein that self-assemble at the cell poles via oligomerization and higher-order assembly, also when ectopically produced in *E. coli* (40–42). Thus, upon induction in *E. coli* cells producing the full-length PopZ protein, the RomR-PopZ^H3H4^-GFP bait fusion protein is targeted to the polar PopZ clusters. In case of direct interaction with RomR, the tested protein fused to mScarlet is recruited to the polar GFP-labelled bait, which can be visualized by fluorescence microscopy. We tested all hits from the RomR proxisome in this assay, except pili-associated proteins which we are currently investigating further. Additionally, we included two known *B. bacteriovorus* RomR partners, i.e., CdgA and SgmX (17, 20). A summary of the results is found in Supp **Fig 5A**. Out of the 11 tested mScarlet-tagged proteins, 7 clearly colocalized with the polar RomR-PopZ^H3H4^-GFP cluster (**Fig 5, Supp Fig 5A-B**). Consistent with previous reports, results from the POLAR assay indicate a direct interaction between RomR and SgmX, but not (or only weakly) between RomR and MglA (20). Moreover, the *B. bacteriovorus* homolog of RomY, which was recently identified in *M. xanthus* as an MglA partner (18), appears as a direct partner of RomR in our assay. Importantly, 5 uncharacterized proteins were recruited by RomR, some of which are predicted to contain domains typically involved in protein-protein interactions, e.g., TPR or coiled-coil domains. These results suggest that RomR interacts directly with a broad range of partners and highlights its role as a polar hub protein.

**FIG 5.**
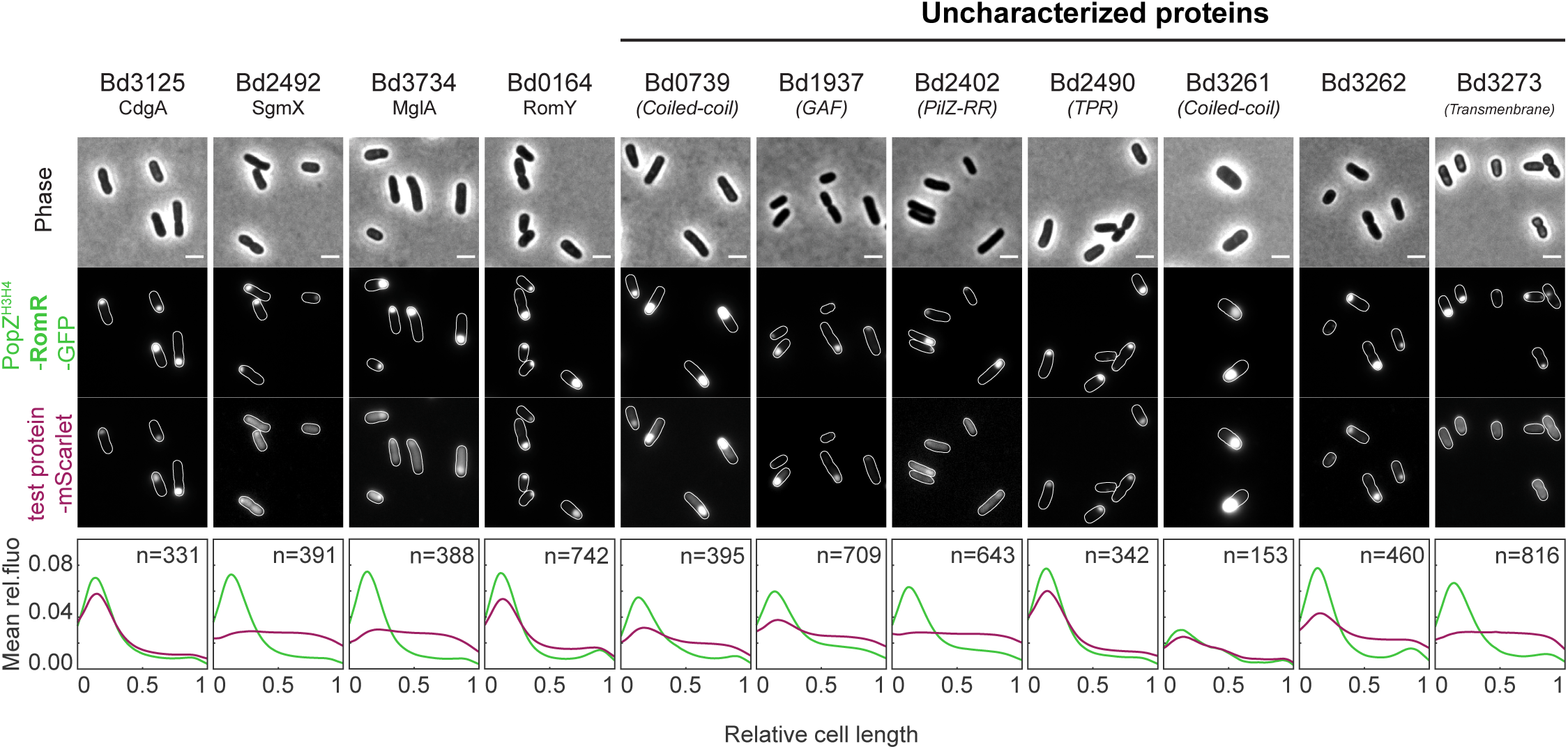
RomR directly interacts with known and uncharacterized proteins of its proximity network. Representative microscopy images of the tested potential RomR partners fused to mScarlet in the presence of the polarly localized PopZ^H3H4^-RomR-GFP (expressed on the “bait vector”). The tested proteins are listed on top with their gene locus. CdgA (Bd3125) and SgmX (Bd2492) were previously reported as RomR partners. MglA (Bd3734) and RomY (Bd0164) are homologues of RomR indirect partners in *M. xanthus.* From top to bottom, the channels are phase contrast, GFP, mCherry. Scale bars are 2 µm. Cell outlines were obtained with Oufti (63). Graphs show the mean pole-to-pole profiles of relative fluorescence intensity in a population of cells, for the PopZ ^H3H4^-RomR-GFP signal in green and the bait-mScarlet fusion in red; “n=” indicates the number of cells analyzed per condition. The control panel with PopZ^H3H4^-GFP (encoded by the “empty” bait vector, i.e., without *romR*) is shown in **Supp Fig 5B**.

### Unlike RomR, Bd0739 and Bd1937 do not form foci during the growth phase

Interestingly, the two top hits from the proximity labelling experiment (Bd0739 and Bd1937) and identified as RomR partners by the POLAR assay are mostly restricted to *B. bacteriovorus* and closely related species within the Bdellovibrionota phylum. Bd0739 is found in a few species, including the epibiotic predator *B. exovorus* but not in *M. xanthus.* Its role is unknown, but the gene is located near *dgcB* (Bd0742), which encodes a diguanylate cyclase proposed to localize at the *B. bacteriovorus* invasive cell pole (21) (**Fig 6A**). When we monitored the localisation of msfGFP-tagged Bd0739 by time-lapse microscopy, a fluorescence signal was only detected when the cells exited the prey (**Fig 6B**, green arrows) where it co-localized with RomR-mCherry expressed in the same cells (**Fig 6B**, pink arrows). Bd1937 is present exclusively in *B. bacteriovorus* but its function is unknown and its close genomic context only encodes hypothetical proteins (**Fig 6C**). The Bd1937-msfGFP signal displayed a pattern reminiscent of Bd0739, as it formed RomR-colocalized spots once the newborn *B. bacteriovorus* progenies left the prey remnants (**Fig 6D**). Our results show that Bd0739 and Bd1937 are proteins of the RomR network that localize at the invasive cell pole upon exit of newborn attack phase predators from the prey. This suggests a role for RomR as a polarity marker, acting as a platform for the localization of proteins related to predatory attack and signalling during the attack phase of *B. bacteriovorus*.

**FIG 6.**
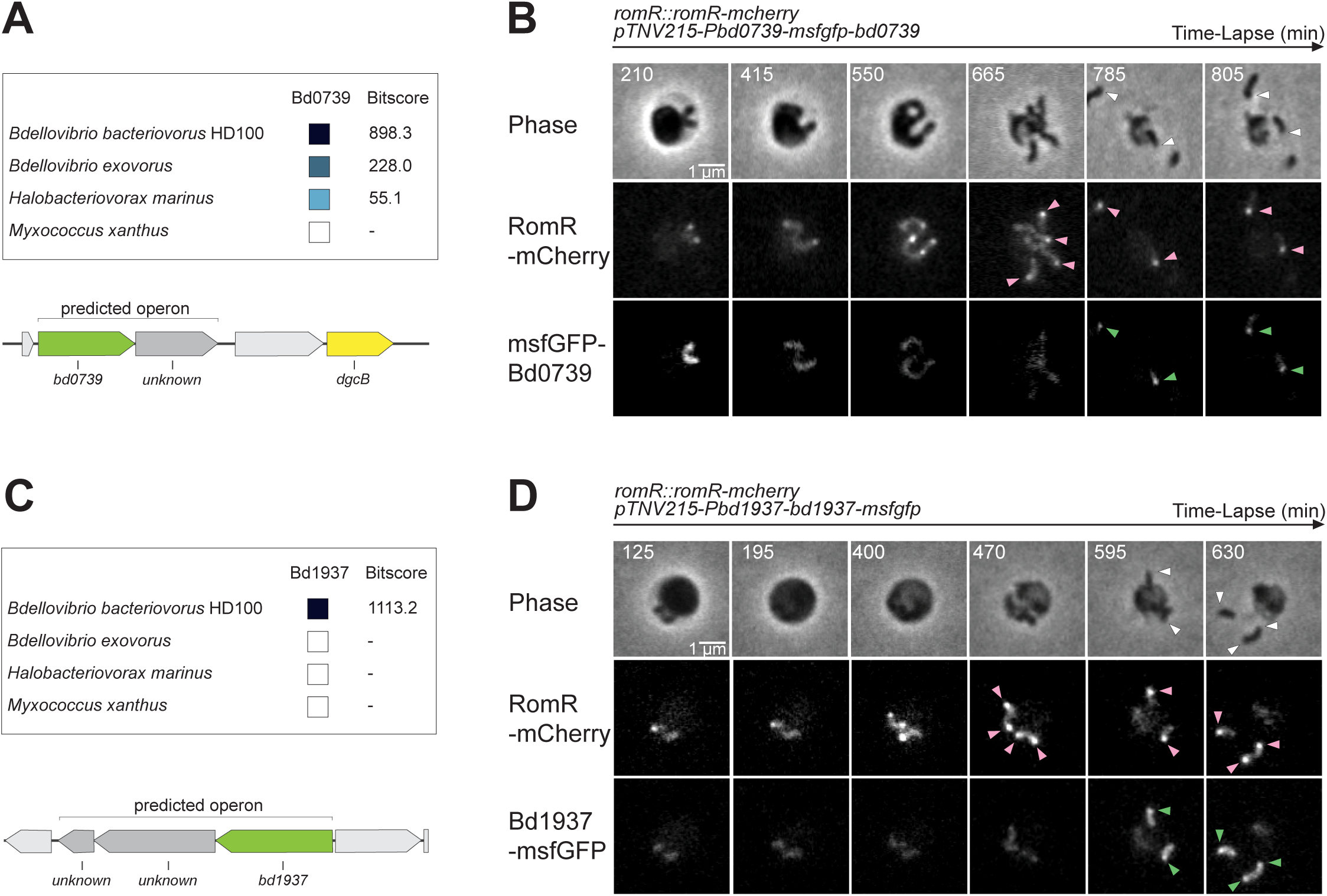
RomR top hit partners Bd0739 and Bd1937 form RomR-colocalized foci in attack phase *B. bacteriovorus* cells. **(A and C)** Schematic representation of the distribution of Bd0739 (A) or Bd1937 (C) homologues in related species: the epibiotic predator *Bdellovibrio exovorus*, another Bdellovibrionota predator *Halobacteriovorax marinus*, and the wolf-pack predator *Myxococcus xanthus*. The bitscores (from BLAST alignments) and corresponding shades of blue were extracted from the String database (https://string-db.org/) using the *B. bacteriovorus* proteins as queries. The genomic context of the protein and operon prediction are illustrated from MicrobesOnline (http://www.microbesonline.org). Unidentified proteins in the predicted operon are coloured in dark grey. The gene *dgcB* is represented in yellow. **(B and D)** Representative time-lapse images of RomR-mCherry and msfGFP-Bd0739 (B, strain GL2381) or Bd1937-msfGFP (D, strain GL2382) during the *B. bacteriovorus* late growth phase and upon release of the predator progeny. Predator cells were mixed with exponentially grown GL655 prey during 90 min prior time-lapse imaging with 5 min intervals. From top to bottom, the channels are phase contrast, mCherry, and GFP. Pink arrowheads point to the new RomR-mCherry foci; green arrowheads to the msfGFP foci (msfGFP-Bd0739 for (B); Bd1937-msfGFP for (D)); white arrowheads to the colocalized foci.

## Discussion

In this study, we provide new insights into the establishment of cell polarity in *Bdellovibrio bacteriovorus*, a bacterium that proliferates through filamentous growth and non-binary division within prey bacteria. Our main findings are summarized in **Figure 7**. First, we found that the RomR protein is an early polarity marker that localizes at the future invasive cell poles of the multiple progenies. Consequently, each daughter cell is equipped with one RomR focus localized at its non-flagellated, invasive pole. Whereas the precise cue that positions RomR remains unknown, our time-lapse monitoring of RomR together with cell cycle progression reporters offers valuable hints into the order of events that take place at the end of the growth phase. RomR foci appear directly at their final locations, *before* cell constriction becomes visible but *after* the appearance of proteins known to be involved in the early assembly stages of the cell division machinery in other species (ZapA and FtsA) (**Figure 7A**). Later, each septum was labeled with a ZapA or FtsA spot, validating fluorescent fusions to these two proteins as relevant early divisome assembly markers in *B. bacteriovorus*. Thus, our results indicate that the labeling of future invasive poles by RomR foci is strictly coordinated in time with the onset of non-binary cell division. However, RomR does not localize at all septa, suggesting the existence of a specific mechanism that facilitates RomR accumulation only at the future invasive cell poles. We previously found that the ParABS system progressively segregates sister chromosomes as they are being asynchronously replicated at different locations in the elongating predator, resulting in an apparent random alternation of *ori-ter* orientations along the filamentous cell (8, 24). Yet, RomR foci appeared after all chromosomal *ori* copies have been synthesized and partitioned in the filamentous predator. Hence, it is tempting to speculate that the spatial organization of the chromosome, in particular the positioning of *ori* copies (which will be located near the invasive cell pole in the progeny), defines the areas between future daughter cells where RomR will accumulate. In this scenario, chromosome orientation would be the earliest determinant of cell polarity in *B. bacteriovorus*. It is interesting to note that the *ori* location impacts the spatial organization of cellular processes in other species, e.g., cell division in *Streptococcus pneumoniae* (43). Regardless of the beacon that attracts RomR at these locations, it is intriguing to note that unlike most described polar hub proteins, RomR in *B. bacteriovorus* finds the right positions along the mother filamentous cell independently of a cell pole.

**FIG 7.**
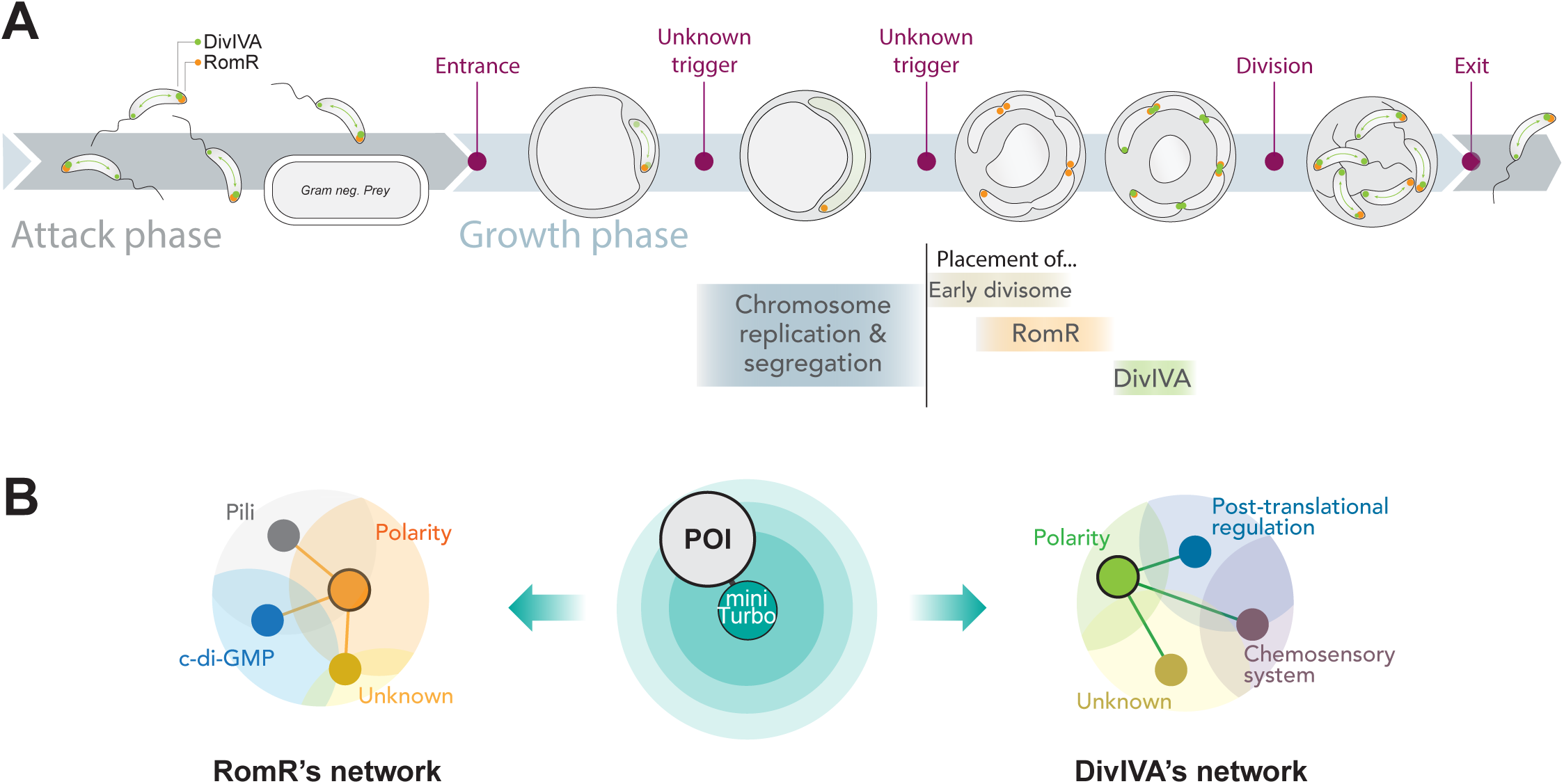
Graphical summary of the *B. bacteriovorus* cell cycle progression (A) and of the proximity labelling results (B). **(A)** The cell cycle of *B. bacteriovorus* is characterized by two main phases: the attack and growth phases, commonly defined based on their presence outside or inside the prey, respectively. The growth phase is characterized by a precise sequence of events visualized with the indicated reporters of key cellular processes, together with RomR and DivIVA. Triggers written in red define distinct periods within the growth phase. The RomR and DivIVA foci are illustrated in orange and green, respectively. **(B)** A protein of interest (POI), coupled with miniTurbo, is illustrated in the centre, while the identified RomR and DivIVA distinct networks are summarized on the left and right, respectively. RomR is represented by an orange circle; DivIVA is represented by a green circle. These circles are connected to other processes via their (in)direct interactions with other proteins.

Extending our investigation of cell polarity in *B. bacteriovorus* to another polar hub candidate, we found that DivIVA and RomR display distinct subcellular localization dynamics during the intra-prey stage of the predatory lifecycle. Unlike RomR, DivIVA became barely detectable when the cells initiated their replicative phase, to reappear only after visible cell constriction started and the establishment of RomR foci at future invasive poles. Furthermore, DivIVA positions itself at all constriction sites and at both “old” poles of the mother predator cell (**Figure 7A**). This pattern likely results from the natural tendency of DivIVA proteins to accumulate at highly curved regions of the cell (10, 11, 32). Yet, DivIVA behaved atypically upon *B. bacteriovorus* cell division and until the start of the proliferative phase inside the prey, switching from pole to pole and alternating between uni- and bipolar localization, while being on average more present at the invasive pole. This strongly suggests that an unknown attack phase-specific mechanism destabilizes the accumulation of DivIVA proteins at curved membranes. Although RomR essentiality prevented us from testing DivIVA’s dependency on RomR for this polar preference in attack phase cells, RomR is unlikely to be responsible for this phenomenon since we did not infer a RomR-DivIVA connection from our proximity labelling assay (discussed below).

Altogether, our time-lapse imaging data highlight that an invariable sequence of events prepares the release and spatial organization of the future offspring at the end of the filamentous growth phase in *B. bacteriovorus*. We recently reported that the time intervals between the final round of chromosome replication and segregation, the end of cell elongation, and the division of the predator cell are remarkably fixed, regardless of the total duration of the growth phase (9). Along the same line, the time intervals between RomR and DivIVA accumulation, chromosome replication and segregation, early divisome assembly, cell constriction, and cell division also seem constant despite the cell-to-cell variability in cell cycle duration (see **Figures 1C, 1E, 3B**). Thus, our new results strengthen the notion that *B. bacteriovorus* proliferation relies on an exquisite spatial and temporal orchestration of different proteins and cellular processes taking place shortly before non-binary cell division (**Figure 7A**).

While the respective functions of RomR and DivIVA remain to be determined, the identification of their close protein network by proximity labelling offers new insights into this question, besides providing supporting evidence for previous models. Despite their shared occupancy at some division sites in the mother *B. bacteriovorus* cell and of the invasive pole of attack phase predators, our proximity labeling approach identified unique protein networks for RomR and DivIVA. This result is consistent with the different localization dynamics of these proteins and emphasizes the specificity of miniTurbo-based proximity labeling for revealing proteins located in the immediate vicinity of the baits. The same approach was recently applied to unveil the protein network of MglA in *M. xanthus*, which included RomR and other proteins of the polarity module (36). Our results provide further support for the idea that RomR connects MglA and cell polarity to key predatory functions via proteins related to Type IV filaments and c-di-GMP signaling (**Figure 7B**). Molecular and functional investigation of these links is ongoing and will be the focus of a future study. It has been proposed and discussed previously by the Sockett group (20, 23) that the triad centered on MglA (RomR-MglA-SgmX), which polarizes pili used for directional motility in *M. xanthus*, has been adapted in *B. bacteriovorus* to polarize pili involved in prey invasion. Our work adds to this model by identifying new proteins in the RomR network, including proteins of unknown function that co-localize with RomR at the invasive cell pole during the attack phase only. We also highlight here the TPR-containing protein Bd2490. BLAST protein sequence alignments show that Bd2490 shares more similarity (18% coverage, e-value 6e-05) with the *M. xanthus* SgmX (MXAN_5766) than Bd2492 (3% coverage, e-value 0.042). We, therefore, propose to rename Bd2492 as SgmX1 and Bd2490 as SgmX2. Both proteins only align with the N-terminal part of *M. xanthus* SgmX involved in Type IV pili activation in *M. xanthus* (44), but not with the C-terminal part required for MglA interaction (45). Their potential role in pili activation in *B. bacteriovorus* and their interaction with MglA remains to be experimentally tested in *B. bacteriovorus*. The finding of several proteins with TPR domains (known to mediate protein-protein interactions) in the RomR proxisome hints at a complex interaction network. Our POLAR experiment suggests that RomR directly interacts with several proteins found in its proximity network, consistent with a polar hub function. Future work will be required to determine the temporality of these interactions during the predatory lifecycle of *B. bacteriovorus*, as it is expected that not all partners bind RomR at the same time.

Whereas a previous study claimed that DivIVA plays a role in *B. bacteriovorus* morphogenesis, our analysis of the Δ*divIVA* mutant did not reveal any cell shape difference compared to the wild-type strain. The reasons for this discrepancy are unclear, but the sequencing of the entire genome of our Δ*divIVA* strain excludes a neutralizing secondary mutation. Besides, the subcellular dynamics of DivIVA, absent during most of the growth phase, might not be compatible with a morphogenesis-related function. Whereas in our hands, cells lacking DivIVA did not present any obvious phenotype, the *in vivo* DivIVA network identified by proximity labelling provides a promising basis for future studies aiming at determining the function of DivIVA. Strikingly, the annotated proteins among the DivIVA proxisome appear to fall within two main categories. A first group of proteins are predicted components of chemosensory pathways (46). While chemotaxis does not seem critical for *B. bacteriovorus* predation (47–49), the particularly large number of chemoreceptors (twenty) encoded in its genome suggests that they serve other signalling functions than chemotaxis (50, 51). These functions may be related to endobiotic proliferation since the epibiotic predator *B. exovorus* only encodes a limited set (three) of chemoreceptors (52). Our results raise the hypothesis that DivIVA might contribute to positioning some of these systems at poles or septa. The DivIVA proxisome also comprises proteins involved in post-translational regulation. Interestingly, the chaperone DnaK might protect DivIVA from degradation in *Staphylococcus aureus* (53), while ClpX, in contrast, is believed to be involved in the degradation of DivIVA in *Streptococcus mutans* (54). Finally, proteins of unknown function were identified in both DivIVA and RomR proxisomes, which might aid their future characterization.

In conclusion, our study offers new insights into the intricate mechanisms governing cell polarity establishment in *B. bacteriovorus* during filamentous growth and non-binary division, revealing RomR as an early polarity marker. Our work underscores the precise spatiotemporal coordination of protein localization and cellular processes leading to the unusual proliferation of this predatory bacterium. Furthermore, the identification of polar protein networks through proximity labelling and POLAR assays strengthens the notion that the invasive cell pole is a specialized platform hosting machineries and pathways critical for predation and implies that DivIVA and RomR represent hubs organizing and connecting different processes in the predator cell. These proximity networks also serve as a foundational step for future investigation into the role of polar proteins and their interacting partners.

## Material and Methods

### Bacterial strains and plasmids

All strains and plasmids used in this study are listed in **Table S1**, and their construction methods are presented in **Table S2**. All oligos used in the study can be found in **Table S3**. Molecular cloning was performed using standard techniques, and DNA assembly was achieved with the NEBuilder HiFi mix (New England Biolabs). The *B. bacteriovorus* strains were obtained from the wild-type HD100 reference strain. The *E. coli* strain used as prey derived from MG1655. All plasmids were introduced into *B. bacteriovorus* by conjugation with the *E. coli* S17-λ*pir* donor strain, following the methodology described previously (24). Scarless allelic replacements into the HD100 chromosome were obtained upon two crossovers of pK18mobsacB-derived suicide vectors. Low-copy replicative plasmids (RSF1010 replicon) were used when indicated “pTNV215”. Chromosomal modifications were screened by PCR and verified by DNA sequencing.

### Routine cultures of B. bacteriovorus and E. coli

For experiments with *E. coli*, overnight starter cultures from single colonies were diluted at least 1:500 in fresh LB medium until exponential phase. For *B. bacteriovorus* cultures, predator cells were grown in DNB medium (Dilute Nutrient Broth, Becton, Dickinson, and Company; supplemented with 2 mM CaCl_2_ and 3 mM MgCl_2_ salts) in the presence of *E. coli* prey at 30°C with continuous shaking as described (55). When using antibiotic-resistant *B. bacteriovorus*, antibiotic-resistant *E. coli* strains were used as prey for the overnight culture. The following antibiotic concentrations were used in liquid and solid media: kanamycin 50 µg/ml, gentamycin 10 µg/ml, ampicillin 50 µg/ml, chloramphenicol 15 µg/ml, tetracycline 7.5 µg/ml.

### Live-cell imaging by phase contrast and epifluorescence microscopy

*B. bacteriovorus* were grown overnight with the appropriate *E. coli* prey, and antibiotics when needed. Prior imaging, *B. bacteriovorus* cells were spotted on 1.2% agarose pads prepared with DNB medium. To stain the flagellum of *B. bacteriovorus* AP cells, FM4-64 stain (Thermo Fisher) was used at a final concentration of 20 µg/ml and cells were incubated in the dark for 2 minutes before imaging. For time-lapse imaging of synchronous predation cycles, unless specified otherwise, the prey MG1655 or GL522 *E. coli* cells were grown in 2TYE medium or LB medium, respectively, until exponential phase (OD_600_ = 0.4-0.6), to obtain larger prey cells as in (9). They were harvested at 5000 x *g* at room temperature for 5 minutes, washed twice, and concentrated in 1/10 initial volume with DNB medium. *E. coli* and *B. bacteriovorus* were then combined in a 1:2 to 1:3 volume ratio, which ensured infection of all *E. coli* cells. In all synchronous predation imaging experiments, the prey-predator mixing step was considered as time 0. The cells were left shaking at 30°C for typically 90 minutes, allowing the predator to invade the prey without phototoxicity. The culture was then deposited on DNB-agarose pads for imaging. At a temperature set to 27°C (Okolab enclosure), several fields of view were repeatedly imaged with intervals of 2.5 to 5 minutes to monitor the growth of *B. bacteriovorus*. For snapshots of *E. coli* cells, except for the POLAR assay (see below), overnight cultures were diluted at least 1:500 with 0.2% arabinose in 1x M9 minimal medium (supplemented with 0.2% glucose, 0.2% casamino acids, 1 µg/ml thiamine, 2 mM MgSO_4_, and 0.1 mM CaCl2). Cells were incubated with shaking at 37°C to reach the exponential phase, and then spotted on 1.2% agarose pads prepared with PBS buffer.

Phase contrast and fluorescence images were taken with a Nikon Ti2-E fully-motorized inverted epifluorescence microscope (Nikon) equipped with a CFI Plan Apochromat l DM 100x 1.45/0.13 mm Ph3 oil objective (Nikon), a Sola SEII FISH illuminator (Lumencor), a Prime95B camera (Photometrics), a temperature-controlled light-protected enclosure (Okolab), and filter-cubes for mCherry (32 mm, excitation 562/40, dichroic 593, emission 640/75; Nikon), and GFP (32 mm, excitation 466/40, dichroic 495, emission 525/50; Nikon). Multi-dimensional image acquisition was managed using the NIS-Ar software (Nikon). By using the 1.5X built-in zoom lens of the Ti2-E microscope, the pixel size was 0.074 µm. Consistent LED illumination and exposure times were employed for capturing images of different strains or conditions within a single experiment.

### POLAR recruitment assay

Protein-protein interaction between the indicated protein pairs was assessed using the POLAR assay (39), following the protocol previously described in (24).

### Prey killing kinetics assay

First, predator cell density was measured for the indicated *B. bacteriovorus* strains in a fresh attack phase population using the SYBR Green labelling assay (55). Then, equal amounts of predators for each condition were mixed with *E. coli* MG1655 in a 96-well microplate, following the procedure described in (55). The plate was shaken continuously at 30°C in a Synergy H1m microplate reader (Biotek) and optical density measurements at 600 nm were taken every 20 minutes. Decrease of OD_600_ indicates prey lysis, as *B. bacteriovorus* cells due to their small size do not affect absorbance readout. Predation kinetics parameters were obtained using the CuRveR package as previously described (55).

### SDS-PAGE and immunoblotting

For immunoblotting, the AP samples were prepared following the method described in (8), starting from 1.5 ml of cleared overnight culture. For the growth phase samples, a 1:1 volume ratio of *E. coli* prey to predator mixture was utilized to minimize the presence of attack phase cells. Samples from the mixed culture were collected every hour and proteins were precipitated with TCA as described in (56). Samples are then loaded onto NuPage 12% Bis-Tris SDS precast polyacrylamide gels (Invitrogen) and run at 190 V for 45 minutes in the NuPAGE MOPS SDS running buffer. Ponceau S staining was applied to the membrane to visualise the total protein, serving as both a loading and transfer control. Standard immunoblotting procedures were performed using a primary antibody against GFP (mouse monoclonal JL-8 antibody, Takara) and a secondary antibody goat anti-mouse IgG-peroxidase antibody (Sigma). Detection of antibody binding was performed by adding luminol to visualize chemiluminescence from the horseradish peroxidase (HRP) reaction and captured with an Image Quant LAS 500 camera (GE Healthcare). Band intensities for the GP samples were quantified using ImageQuantTL software, normalized by the Ponceau S staining. The results were graphically presented using GraphPad Prism.

For the immunoblots before the preparation of proximity labelling samples (i.e., to evaluate the endogenous biotinylation of *B. bacteriovorus* and the distinct biotinylation pattern of each bait (**Supp Fig 4A**)), the protocol was followed as described above, using Streptavidin-HRP (Fisher Scientific).

For the analysis of proximity labelling samples (see text below, and **Supp Fig 4B**), sample loading buffer (Bio-Rad XT sample buffer) and reducing agent (Bio-Rad) was added to the samples according to the manufacturer’s instruction and equivalent amounts of protein (∼5 µg) were separated on a 4 to 12% gradient XT precast Criterion gel using XT-MOPS buffer (1.0 mm thick 4 to 12% polyacrylamide Criterion Bis-Tris XT-gels, Bio-Rad) at 150–200V. Subsequently, proteins were transferred onto a PVDF membrane. Membranes were blocked for 30 min in a 1:1 Tris-buffered saline (TBS)/Odyssey blocking solution (cat n° 927-40003, LI-COR, Lincoln, NE, USA) and probed by immunoblotting. Following overnight incubation of primary antibody in TBS-T/Odyssey blocking buffer and three 10 min washes in TBS-T (0.1% Tween-20), membranes were incubated with secondary antibody for 30 min in TBS-T/Odyssey blocking buffer followed by 3 washes in TBS-T or TBS (last washing step). The following antibodies or detection reagents were used: mouse anti-Flag (Sigma, F3165; 1/4000), streptavidin-Alexa Fluor™ 680 Conjugate (Invitrogen, S32358, 1/10000) and anti-mouse (IRDye 800 CW goat anti-mouse antibody IgG, LI-COR, cat n° 926-32210, 1/10000) and bands were visualized using an Odyssey infrared imaging system (LI-COR).

### Proximity labelling with miniTurbo

The level of endogenous biotinylation in predator and prey was first tested to estimate the quantity of biotinylated proteins and thus the feasibility of using proximity labelling in our *B. bacteriovorus*. Samples of *E. coli* and *B. bacteriovorus* in AP and GP were used for immunoblotting with Streptavidin, which showed only a few biotinylated bands and the absence of smear). The toxicity of various concentrations of biotin (0 - 4 mM) was also assessed by killing assay as described above. Biotin concentrations from 0 to 1 mM did not impact the *B. bacteriovorus* predatory lifecycle. Given the growth of *B. bacteriovorus* within another bacterium, several optimizations were carried out, including incubation time, biotin concentration, and incubation of prey with biotin prior to predation. The optimized workflow used in this study is as follows: starting with a cleared overnight culture in DNB medium of *B. bacteriovorus*, the quantity of predators for each condition is assessed by SYBR Green (55). The culture is then supplemented with 50 µM biotin, from a 50 mM biotin stock solution prepared in water with 60 mM NaOH (lyophilized biotin powder, 500 mg stock, Merck Life Science BV (Sigma)). The biotin-supplemented culture was then incubated for 2 h at 30°C with shaking, without prey to allow labelling of proximal proteins (“proxisome”) in attack phase *B. bacteriovorus*. For proximity labelling of growth phase *B. bacteriovorus* cells, attack phase predators were incubated with biotin as described above, and wild-type *E. coli* cells were also incubated with biotin to maximize the biotin supply to *B. bacteriovorus* within its prey. Briefly, from an overnight culture from a colony in LB medium, a 100-foled diluted *E. coli* culture was prepared in fresh medium supplemented with 50 µM biotin. The biotin-supplemented prey culture was placed in a shaking incubator at 37°C for 2 h before washing in DNB medium to serve as a prey suspension. Following the 2 h biotin incubation of the predators, the *B. bacteriovorus* and prey cultures (at OD_600_ = 0.5) were used to inoculate into fresh DNB medium, supplemented with biotin to maintain a final concentration of 50 µM. This was followed by 5 h incubation at 30°C with shaking to allow the predators to complete their growth phase. Cells were harvested by centrifugation of the predation mix at 7000 x g for 10 min, then frozen or used directly for immunoblot analysis to assess the biotinylation profiles specific to each construct (as seen in **Supp Fig 4A**).

Preparation of proximity labelling samples for MS was essentially performed as described in (57). More specifically, protein extraction was performed by resuspension of bacterial pellets in lysis buffer (100 mM Tris pH 7.5, 8 M urea, 150 mM NaCl, 2% SDS), followed by three rounds of freeze-thawing in liquid nitrogen, and microprobe sonication. After centrifugation, a fraction of the lysates was taken as a control for immunoblot (**Supp Fig 4B**), the rest was resuspended with streptavidin-coated beads (Streptavidin Sepharose High Performance, # GE17-5113-01, Sigma-Aldrich). After overnight incubation in a rotator at room temperature (RT), the supernatant was removed (and used as the “unbound fraction” for immunoblotting; **Supp Fig 4B**) and 4 series of washes (by repeated resuspension and pelleting of the beads at 600 x g for 5 min) were performed at RT for 5 min followed by a 30 min wash in high salt buffer (100 mM Tris-HCL pH7.5, 1M NaCl) and finally resuspend in ultrapure HPLC (high performance liquid chromatography) grade water. A small fraction (1/10) was used for immunoblotting (“bound fraction”, concentrated 5x in elution buffer (2% SDS, 3 mM biotin, 8M urea, PBS; **Supp Fig 4B**)) and the rest was digested by trypsin after resuspension of the beads in ABC buffer (50 mM ammonium bicarbonate pH 8.0, 1.0 µg trypsin per sample) and overnight incubation at 37°C with agitation. After overnight digestion, an additional 0.5 µg of trypsin was added, followed by an additional 2-hours incubation with shaking at 37°C. The beads were pelleted (600 x g, 3 min) and the supernatant transferred to a fresh low protein binding tube (Eppendorf). The beads were washed with 1 x 200 µL HPLC-grade water and the washes combined with the original supernatant. The peptide solution was acidified with 5% TFA (trifluoroacetic acid) to reach a final concentration of 0.1%, cleared from insoluble particles by centrifugation for 10 min at 16,000 x g (4°C), and the supernatant transferred to clear tubes. The samples were dried in a SpeedVac concentrator and were resuspended in 100 µl of 0.1% TFA. Methionine oxidation was performed by the addition of H_2_O_2_ to reach a final concentration of 0.5 % (v/v) for 30 minutes at 30°C. Solid phase extraction of peptides was done using equilibrated C18 pipette tips (Bond Elut OMIX 100 µl pipette tips, Agilent). The samples are aspirated for 10 cycles for maximum binding efficiency and then eluted in LC-MS/MS vials with 100 µl of 0.1% TFA in water:acetonitrile (at a ratio of 30:70 (v/v)). The bound peptides were finally dried in a SpeedVac concentrator and dissolved in a resuspension buffer (composed of 2 mM Tris(2-carboxyethyl)phosphine (TCEP) in water: LC-grade acetonitrile (ACN) (98:2 (v/v)).

### LC-MS/MS and data analysis

Per bait, 3 replicate samples were analysed by LC-MS/MS using an UltiMate 3000 RSLC nano HPLC (Dionex) in-line connected to a Q Exactive HF mass spectrometer (Thermo Fisher Scientific Inc.) equipped with a Nanospray Flex Ion source (Thermo) as previously described (58, 59). Raw data files were searched with MaxQuant (60) using the Andromeda search engine (61) (version 1.6.10.43) and MS/MS spectra searched against the UniProt UP000008080 database (taxonomy *Bdellovibrio bacteriovorus* (strain ATCC 15356 / DSM 50701 / NCIMB 9529 / HD100)) (version 2022_11, containing 3583 protein entries; https://www.uniprot.org/proteomes/UP000008080)) based on the GCA_000196175.1 genome assembly and annotation from ENA/EMBL and complemented with the miniTurbo-Flag sequence. Potential contaminants present in the contaminants.fasta file that comes with MaxQuant were automatically added. A precursor mass tolerance was set to 20 ppm for the first search (used for nonlinear mass recalibration) and set to 4.5 ppm for the main search. As enzyme specificity, trypsin was selected and one missed cleavages was allowed. Methionine oxidation was set as fixed modifications and no variable modifications were specified. The false discovery rate (FDR) for peptide and protein identification was set to 1%, and the minimum peptide length was set to 7. The minimum score threshold for both modified and unmodified peptides was set to 40. The match between runs function was disabled and proteins were quantified by the MaxLFQ algorithm integrated in the MaxQuant software (62). Here, a minimum of two ratio counts and only unique peptides were considered for protein quantification.

For basic data handling, normalization, statistics and annotation enrichment analysis, we used the freely available open-source bioinformatics platform Perseus (version 1.6.15.0) (Tyanova, Temu et al. 2016). Data analysis after uploading the protein groups file obtained from MaxQuant database searching was performed as described previously (59). In brief, all replicate samples were grouped and LFQ-intensities log_2_ transformed. Reference groups were defined as the complement group of each setup (i.e., the union of all other setups). Proteins with less than three valid values in at least one group were removed and missing values were imputed from a normal distribution around the detection limit (with 0.3 spread and 1.8 down-shift). Then, a *t* testing (FDR= 0.01, S0 = 0.1) was performed to detect enrichments in the different setups (**Fig 4B-C, Supp Fig 4C-E**). For more information, see the detailed protocol in (57).

### Image analysis

The software Oufti (63) was used to detect cell outlines and diffraction-limited fluorescent foci of AP *B. bacteriovorus* cells with subpixel precision from, respectively, phase contrast images and fluorescence images. The detected spots were attributed to the corresponding cell in the Oufti cell lists, which includes coordinates, morphology, and signal intensity data. Histograms of foci numbers per cell and distributions of morphological parameters (length, width, curvature) were then plotted using custom MATLAB (Mathworks) codes as in (8). Demographs of relative fluorescence intensity in cells sorted by length were plotted as in (8, 64). Arrays of relative fluorescence intensity values were oriented based on the position of the maximal fluorescence intensity of the indicated signal in each cell half. Kymographs were obtained using the built-in kymograph function in Oufti (63). Killing curves were generated using the CurveR package (55). Graphs of the mean relative fluorescence profiles (for POLAR assay) were plotted using the custom MeanIntProfile MATLAB code as in (8). The cells were oriented based on the maximal fluorescence intensity of the PopZ^H3H4^-(RomR-)GFP. For the graphs in **Fig 1C, 1E, 3B**, the frame numbers of new foci appearance were reported in an Excel table for each bdelloplast and plotted using GraphPad Prism.

### Image processing

Images were processed using FIJI (65), applying identical contrast and brightness settings per channel for all figures from the same experiment. Since time-lapses images were taken with low exposure to avoid phototoxicity, denoising (Denoise.ai, Nikon) was applied for display purposes only. Time-lapse image stacks were corrected for small drifts between frames using the ImageStabilizer plugin in FIJI (66)), applied to all channels. *B. bacteriovorus* outlines were drawn manually in Adobe Illustrator for AP and GP cells within bdelloplasts, unless stated otherwise. *E. coli* outlines were obtained using Oufti (63). Microscopy images, graphs, outlines, illustrative figures were assembled and labelled using Adobe Illustrator.

## Legends or the supplementary figures

**SUPP FIG 1** Characterization of the localisation of RomR during the *B. bacteriovorus* growth phase. **(A)** Representative bdelloplast containing a GP *B. bacteriovorus* cell of the *romR::romR-mcherry* strain (GL1466) in which no second polar RomR-mCherry focus is visible during the growth of the filament. New foci appear along the cell body. The predators were mixed with exponentially grown MG1655 *E.coli* prey for 90 min prior time-lapse imaging with 5 min intervals. The phase contrast channel is on top, the mCherry channel at the bottom. Dark pink arrowheads point to the “old” RomR-mCherry focus at the invasive pole, light pink arrowheads to the new foci appearing during the growth phase, and asterisks to constriction sites. **(B)** Graphical representation of the percentage of cells with a unipolar (53,3%; as in A) or bipolar (46,7%, as in **Fig 1A**) localisation of RomR-mCherry foci during growth (n=30). **(C)** Localisation of RomR-mCherry compared to the early divisome component msfGFP-FtsA (GL2379). Representative bdelloplast containing a GP *B. bacteriovorus* cell (strain GL2379) mixed with exponentially grown GL522 *E. coli* prey for 90 min prior time-lapse imaging with 5 min intervals. From top to bottom, phase contrast, mCherry, GFP channels and overlay of both FP channels with RomR-mCherry in magenta and msfGFP-FtsA in green. The dark pink arrowhead points to the old RomR-mCherry focus, light pink arrowheads point to the new RomR-mCherry foci; green arrowheads to the msfGFP-FtsA foci; white arrowheads to the colocalized foci. Each time-lapse experiment was performed at least 3 times. *B. bacteriovorus* filament and bdelloplasts outlines were drawn manually based on the phase contrast images.

**SUPP FIG 2** Characterization of the localisation of DivIVA during the *B. bacteriovorus* attack and growth phases. **(A)** DivIVA-msfGFP localizes preferentially at the non-flagellated pole, visualized using the origin of replication on the chromosome (*ori*) via the *parS_PMT1_·*YFP-ParB_PMT1_ system previously used in *B. bacteriovorus* (8). Left: Schematic representation of an attack phase cell with the nucleoid in blue and the *ori* as a blue dot near the invasive pole. Right: Demographs of the fluorescent signals imaged in attack phase GL1636 cells (YFP-ParB _PMT1_ on the left, DivIVA-msfGFP on the right), sorted by length and oriented with the YFP-ParB_PMT1_ (*ori* marker) signal on the left, labelled as pole 1. Blue-to-red indicates low-to-high fluorescence intensities; the number of analysed cells is (n=)1773. **(B)** DivIVA_Bb_-msfGFP uniformly localizes at both poles and septa when heterologously expressed in a *E. coli*. Left: Representative images of MG1655 *E. coli* strain expressing DivIVA-msfGFP from an inducible promoter on a plasmid (pBAD18). Phase channel on the left, GFP channel on the right. Right: Demograph of the DivIVA-msfGFP fluorescent signal in *E. coli*, sorted by length, with random orientation. White-to-red indicates low-to-high intensities; the number of analysed cells is (n=)2028. **(C)** DivIVA-msfGFP protein amount during the *B. bacteriovorus* cell cycle. A clear culture of AP cells for the *B. bacteriovorus divIVA::divIVA-msfgfp* strain (GL1620) was mixed with stationary phase MG1655 *E. coli* prey and samples from this predation mix were collected every hour and used for detection of DivIVA-msfGFP by immunoblot (C) or imaged (D). DivIVA-msfGFP protein amount increases at the end of the cell cycle. Left: whole-cell protein extracts from the time-course samples were used for a immunoblotting with an anti-GFP antibody. Attack and growth phase samples were loaded on the same gel but only GP samples are compared since the loading material differs between the two phases. Molecular weight markers (kDa) are shown on the side. Ponceau staining (bottom) was performed to serve as a loading and transfer control and thus to be used for normalisation as done previously (24). Right: Quantification of the relative protein level for each GP timepoint, normalised by the corresponding Ponceau lane. **(D)** Localisation of DivIVA-msfGFP in *B. bacteriovorus* growing in smaller prey cells. Left: Representative phase contrast and GFP channel images of AP cells for the *B. bacteriovorus divIVA::divIVA-msfgfp* strain (GL1620). Right: Representative bdelloplasts containing a GP cell of the same strain imaged in time-course upon mixing with *E. coli* MG1655 as in C. Scale bars are 1 µm. *B. bacteriovorus* cells and bdelloplasts outlines were drawn manually based on the phase contrast images.

**SUPP FIG 3** The absence of DivIVA does not impact the morphology of *B. bacteriovorus* (AP) cells. **(A)** Representative phase contrast images of the wild-type HD100 strain (GL734, left) and the HD100 Δ*divIVA* strain (GL1641, right). **(B)** Histograms of cell width, curvature, and cell length, comparing the AP cells of the wild-type and Δ*divIVA* strains. Mean values and graphical elements related to wild-type and Δ*divIVA* strains are coloured in blue and purple, respectively. The number of cells analysed is (n=) 881 for wild-type and 1812 for Δ*divIVA*. **(C)** Killing curves showing the percentage of prey survival over time when the *E. coli* prey are mixed with *B. bacteriovorus* wild-type (blue) or *ΔdivIVA* (purple) strains. The maximum killing rate is given by the “r_max_” values and the time when this maximum rate is obtained is given by the “s=” values. Mean curves obtained from three biological replicates are shown as plain or dotted lines, respectively. Values were normalized to the initial absorbance value.

**SUPP FIG 4** Endogenous miniTurbo-based proximity labelling in *B. bacteriovorus*. (8) **(A)** Immunoblot using a Streptavidin-HRP conjugate on whole-cell protein extract to evaluate the distinct biotinylation pattern of each bait (fused to miniTurbo and expressed from their native locus). Biotin (50 µM) was added for 2h during the attack phase followed by 5h during the growth phase. Molecular weight markers (kDa) are shown on the side. Distinct biotinylation patterns indicate POI-specificity and *cis-* and *trans-*biotinylation activity of the protein fusions. **(B)** Immunoblots of fractions collected during the sample preparation for MS, demonstrating the efficient and specific capture of biotinylated protein material. Top: An anti-Flag antibody was used for the detection of the indicated bait protein fusions to the miniTurbo-Flag enzyme. Middle: A streptavidin/Alexa FluorTM 680 (S680) conjugate was used for the detection of biotinylated proteins. Bottom: Overlay of both immunoblots (anti-Flag in green and S680 in red). The overlay (yellow) signal shows that the bait fusion is biotinylated (as also pointed by the arrowheads on the two top blots). Unspecific signal (asterisks) and endogenously biotinylated protein (star) serve as loading controls. For each bait, the input samples of 3 replicates were analysed (I1 to 3), alongside the streptavidin unbound (U) and bound (B) fractions (5x concentrated) of the first replicate sample. **(C)** Heat map visualization of pairwise LFQ (label free quantification) correlations (Pearson), averaged per setup across the replicates (3 for each bait) analysed by MS following on-bead tryptic digestion, indicating a good correlation among the replicates while showing the variability between the replicates groups (i.e., the different baits). **(D)** Heat map representation of cluster analysis after ANOVA. The intensities of proteins with significantly different abundance (p ≤ 0.01) in the respective proxisomes (total protein #100) are represented. Three clusters (clusters 1-3), respectively nearly exclusively enriched for RomR (Cluster 2), DivIVA (Cluster 3) and ParB (Cluster 1) putative interactors can be observed. Green-to-red indicates low-to-high intensities.

**SUPP FIG 5** RomR directly interacts with several proteins of its proximity network. **(A)** Schematic representation of the POLAR assay results using RomR as a bait. The proteins used as prey are listed on top with their gene locus. The first row corresponds to the results obtained with the empty bait vector (shown in **B**), containing only PopZ^H3H4^-GFP, while the second row corresponds to the results obtained with RomR as a bait (shown in **Fig 5**). Localization patterns observed for the mScarlet-tagged protein, with or without the RomR bait, are color-coded as shown on the right. We conclude on RomR-dependent recruitment, indicating direct interaction, for proteins in the light green (diffuse and foci) and dark green (foci) categories in the presence of RomR. **(B)** Representative microscopy images of the tested potential RomR partners tagged with mScarlet in the presence of the polarly localized PopZ^H3H4^-GFP (encoded by the “empty” vector without bait). From top to bottom, the channels are phase contrast, GFP, mCherry. Scale bars are 2 µm. Cell outlines were obtained with Oufti (63). Graphs show the mean pole-to-pole profiles of relative fluorescence intensity in a population of cells, for the PopZ ^H3H4^-GFP signal in green and the bait-mScarlet fusion in red; “n=” indicates the number of cells analyzed per condition.

## Acknowledgements

We are grateful to Charles de Pierpont for excellent technical support, Terrens Saaki for the construction of ZapA-msfGFP and msfGFP-FtsA fusions, members of the Laloux lab for insightful discussions and critical reading of the manuscript. This work was supported by the European Commission (ERC Starting grants PREDATOR 802331 to G.L. and PROPHECY 803972 to P.V.D.), the F.R.S.-FNRS (Mandat d’Impulsion Scientifique MIS-F.4521.18 MICRO-PREDATOR to G.L.), the Fonds Jacques Moulaert (Fondation Roi Baudoin) to G.L., and the Research Foundation Flanders (FWO-Vlaanderen) (project numbers G051120N and G045921N, to P.V.D.). Y.S. is a post-doctoral fellow of the European Molecular Biology Organization, C.T. is a Research Fellow of the F.R.S.-FNRS, J.K. was a Research Fellow of the F.R.S.-FNRS, G.L. is a Research Associate of the F.R.S.-FNRS.

