## Supplementary Tables 1 to 3 for "Distinct dynamics and proximity networks of hub proteins at the prey-invading cell pole in a predatory bacterium"

#### Table S1 – Strains used for this study

| Strains | Description | Resistance | Source |
|---|---|---|---|
| ***Bdellovibrio bacteriovorus*** | | | |
| **GL734** | Wild-type *B. bacteriovorus* HD100 | - | Lab collection (Kind gift from R.E. Sockett, U. Nottingham) |
| **GL806** | HD100 *ori*::*parS*$_{PMT1}$ | - | [1] |
| **GL944** | HD100 *divIVA::divIVA-msfgfp* / pTNV215-*romR-tdtomato* | Gm | This study |
| **GL1466** | HD100 *romR::romR-mcherry* | - | This study |
| **GL1471** | HD100 *divIVA::divIVA-msfgfp romR::romR-mcherry* | - | This study |
| **GL1620** | HD100 *divIVA::divIVA-msfgfp* | - | This study |
| **GL1636** | HD100 *divIVA::divIVA-msfgfp ori::parS*$_{PMT1}$ / pTNV215-*mcherry-parB*$_{PMT1}$ | Gm | This study |
| **GL1641** | HD100 Δ*divIVA* | - | This study |
| **GL1655** | HD100 *romR::romR-mcherry parB::parB-yfp* | - | This study |
| **GL1941** | HD100 *parB::parB-miniTurbo-flag* | - | This study |
| **GL1985** | HD100 *romR::romR-miniTurbo-flag* | - | This study |
| **GL1988** | HD100 *divIVA::divIVA-miniTurbo-flag* | - | This study |
| **GL2378** | HD100 *romR::romR-mcherry* / pSEVA251-*PzapA-zapA-msfgfp* | Kan | This study |
| **GL2379** | HD100 *romR::romR-mcherry* / pSEVA251-*PftsA-msfgfp-ftsA* | Kan | This study |
| **GL2380** | HD100 Δ*divIVA romR::romR-mcherry* | - | This study |
| **GL2381** | HD100 *romR::romR-mcherry* / pTNV215-*msfgfp-bd0739* | Gm | This study |
| **GL2382** | HD100 *romR::romR-mcherry* / pTNV215-*bd1937-msfgfp* | Gm | This study |
| ***Escherichia coli*** | | | |
| **TOP10** | Strain used for bait plasmid cloning (POLAR assay) | - | Lab collection |
| **CC118 λpir** | Strain used for prey plasmid cloning (POLAR assay) | - | Kind gift from T. Bernhardt[2] |
| **TB28/pAH69** | Strain used for the double transformation (POLAR assay) | Amp | Kind gift from T. Bernardt[2] |
| **GL58** | MC4100 / pBAD18 | Amp | Lab collection |
| **GL522** | MG1655 *ΔmreB* / pRM-*mreB(L251R)CD*, used as prey | Chlor | Kind gift from K.C. Huang[3], described as prey for *B. bacteriovorus* in [4] |
| **GL606** | NEB5α / pTNV215-*tdtomato* | Gm | [1] |
| **GL611** | S17-1 *λpir* / pTNV215-*romR-tdtomato* | Gm | [1] |
| **GL633** | S17-1 *λpir* / pK18mobsacB-*bd0464up-divIVA-msfgfp-bd0464down* | Kan | This study |
| **GL655** | Wild-type *E. coli* MG1655, used as prey | - | Lab collection |
| **GL669** | TOP10 / pK18mobsacB | Kan | Lab collection |
| **GL728** | DH5α / pBG18 (pSEVA251-*pBioFab-sfgfp*) | Kan | Kind gift from S. Bigot & C. Lesterlin |
| **GL891** | Strain containing the vector pHCL149 (POLAR assay) | Tet | Kind gift from T. Bernhardt[2] |

| | | | |
|---|---|---|---|
| **GL893** | Strain containing the vector pHCL150 (POLAR assay) | Chlor | Kind gift from T. Bernhardt[2] |
| **GL894** | Strain containing the vector pHCL151 (POLAR assay) | Tet | Kind gift from T. Bernhardt[2] |
| **GL983** | S17-1 λ*pir* / pK18mobsacB-*bd3907up-parB-yfp-bd3907down* | Kan | This study |
| **GL1001** | S17-1 λ*pir* / pTNV215-*mcherry-parB*$_{PMT1}$ | Gm | [1] |
| **GL1038** | MG1655 / pBAD18 *divIVA-msfgfp* | Amp | This study |
| **GL1468** | TOP10 / pENTR_R2_L3_GGGGS_miniTurbo_FLAG | Kan | Petra Van Damme lab collection |
| **GL1638** | S17-1 λ*pir* / pK18mobsacB-*bd2761up-romR-mcherry-bd2761down* | Kan | [5] |
| **GL1640** | S17-1 λ*pir* / pK18mobsacB-*bd0464up-bd0464down* | Kan | This study |
| **GL1909** | S17-1 λ*pir* / pK18mobsacB-*bd3907up-parB-miniTurbo-flag-bd3907down* | Kan | This study |
| **GL1984** | S17-1 λ*pir* / pK18mobsacB-*bd2761up-romR-miniTurbo-flag-bd2761down* | Kan | This study |
| **GL1987** | S17-1 λ*pir* / pK18mobsacB-*bd0464up-divIVA-miniTurbo-flag-bd0464down* | Kan | This study |
| **GL2301** | TOP10 / pHCL150-*romR* as Bait for POLAR assay | Chlor | This study |
| **GL2384** | CC118 λ*pir* / pHCL151-*mscarlet-bd0739* | Tet | This study |
| **GL2385** | CC118 λ*pir* / pHCL147-*bd1937-mscarlet* | Tet | This study |
| **GL2386** | CC118 λ*pir* / pHCL147-*bd2402-mscarlet* | Tet | This study |
| **GL2388** | CC118 λ*pir* / pHCL147-*bd3734-mscarlet* | Tet | This study |
| **GL2390** | CC118 λ*pir* / pHCL147-*bd2490-mscarlet* | Tet | This study |
| **GL2391** | CC118 λ*pir* / pHCL147-*bd2492-mscarlet* | Tet | This study |
| **GL2392** | CC118 λ*pir* / pHCL147-*bd3125-mscarlet* | Tet | This study |
| **GL2394** | CC118 λ*pir* / pHCL147-*bd3262-mscarlet* | Tet | This study |
| **GL2395** | CC118 λ*pir* / pHCL151-*mscarlet-bd3261* | Tet | This study |
| **GL2396** | CC118 λ*pir* / pHCL151-*mscarlet-bd3273* | Tet | This study |
| **GL2397** | CC118 λ*pir* / pHCL147-*bd0164-mscarlet* | Tet | This study |

#### Table S2 – Strains and plasmids construction

| **Strains** | **Construction** |
|---|---|
| **GL944** | Mating GL1620 (HD100) x GL611 |
| **GL1038** | Transformation GL655 (MG1655) x GL1034 miniprep |
| **GL1466** | Mating GL734 (HD100) x GL1638, allelic replacement |
| **GL1471** | Mating GL1620 (HD100) x GL1638, allelic replacement |
| **GL1620** | Mating GL734 (HD100) x GL633, allelic replacement |
| **GL1636** | Mating GL806 (HD100) x GL633, allelic replacement, followed by mating of the resulting strain x GL1001 |
| **GL1641** | Mating GL734 (HD100) x GL1640, allelic replacement |
| **GL1655** | Mating GL1466 (HD100) x GL983, allelic replacement |
| **GL1941** | Mating GL734 (HD100) x GL1909, allelic replacement |
| **GL1985** | Mating GL734 (HD100) x GL1984, allelic replacement |
| **GL1988** | Mating GL734 (HD100) x GL1987, allelic replacement |
| **GL2378** | Mating GL1466 (HD100) x GL1162 |
| **GL2379** | Mating GL1466 (HD100) x GL1161 |
| **GL2380** | Mating GL1466 (HD100) x GL1640, allelic replacement |
| **GL2381** | Mating GL1466 (HD100) x GL2398 |
| **GL2382** | Mating GL1466 (HD100) x GL2399 |
| **Plasmid** | **Construction** |
| GL633: pK18mobsacB-*bd0464up-divIVA-msfgfp-bd0464down* | Assembly of the following PCR-amplified fragments: opened vector pK18mobsacB from GL669 miniprep with primers oGL331-332; *bd0464up* (including *divIVA [bd0464]*) from HD100 gDNA with primers oGL333-334; *4GS_linker-msfgfp* with oGL299-337; and *bd0464down* from HD100 gDNA with primers oGL335-336. |
| GL983: pK18mobsacB-*bd3907up-parB-yfp-bd3907down* | Assembly of the following PCR-amplified fragments: opened vector pK18mobsacB from GL669 miniprep with primers oGL331-332; *bd3907up* (including *parB [bd3907]*) from HD100 gDNA with primers oGL751-752; *4GS-yfp* with primers oGL286-287; and *bd03907down* from HD100 gDNA with primers oGL754-288. |
| GL1034: pBAD18-*divIVA-msfgfp* | Assembly of the following PCR-amplified fragments: opened vector pBAD18 from GL58 miniprep with primers oGL991-992; and *divIVA-4GS-msfgfp* from GL633 with primers oGL993-994. |
| GL1161: pSEVA251-*Promoter(ftsA)-msfgfp-ftsA* | Assembly of the following PCR-amplified fragments: opened vector pSEVA251 from GL728 miniprep with primers oGL897-677; *PftsA* from HD100 gDNA with primers oGL931-664; and *msfgfp-4GS-ftsA(bd3190)* with primers oGL661-932 from GL1272 (TOP10 / pBAD18-*msfgfp-ftsA*, constructed using the opened vector pBAD18 from GL58 miniprep with primers oGL264-322, *msfgfp-4GS* with primers oGL399-400, *ftsA* from HD100 gDNA with primers oGL553-554). |
| GL1162: pSEVA251-*Promoter(zapA)-zapA-msfgfp* | Assembly of the following PCR-amplified fragments: opened vector pSEVA251 from GL728 miniprep with primers oGL897-677; *PzapA* (including *zapA [bd1185]*) from HD100 gDNA with primers oGL929-308; and *4GS-msfgfp* with primers oGL299-930. |
| GL1640: pK18mobsacB-*bd0464up-bd0464down* | Assembly of the following PCR-amplified fragments: opened vector pK18mobsacB from GL669 miniprep with primers oGL502-265; *bd0464up* (with the 2 first codons of *divIVA [bd0464]*) from HD100 gDNA with primers oGL503-1211; and *bd0464down* (with the 3 last codons and stop codon of *divIVA*) from HD100 gDNA with primers oGL1212-508. |
| GL1909: pK18mobsacB-*bd3907up-parB-miniTurbo-flag-bd3907down* | Assembly of the following PCR-amplified fragments: opened vector pK18mobsacB from GL669 miniprep with primers oGL331-332; *bd3907up* (including *parB [bd3907]*) from HD100 gDNA with primers oGL752-751; *4GS-miniTurbo-flag* from GL1468 miniprep with primers oGL1767-1768; and *bd3907down* from HD100 gDNA with primers oGL754-1769. |

| | |
|---|---|
| GL1984: pK18mobsacB-*bd2761up-romR-miniTurbo-flag-bd2761down* | Assembly of the following PCR-amplified fragments: opened vector pK18mobsacB from GL669 miniprep with primers oGL784-332; *bd2761up* (including *romR [bd2761]-4GS*) from HD100 gDNA with primers oGL1236-1628; *miniTurbo-flag* from GL1468 miniprep with primers oGL1629-1729; and *bd2761down* from HD100 gDNA with primers oGL1730-1238. |
| GL1987: pK18mobsacB-*bd0464up-divIVA-miniTurbo-flag-bd0464down* | Assembly of the following PCR-amplified fragments: opened vector pK18mobsacB from GL669 miniprep with primers oGL331-332; *bd0464up* (including *divIVA [bd0464]-4GS*) from HD100 gDNA with primers oGL334-1633; *miniTurbo-flag* from GL1468 miniprep with primers oGL1629-1729; and *bd2761down* from HD100 gDNA with primers oGL1731-335. |
| GL2301: pHCL150-*romR* | Assembly of the following PCR-amplified fragments: opened vector pHCL150 from GL893 with primers oGL2464-2465; and *romR* from HD100 gDNA with primers oGL2485-2486. |
| GL2384: pHCL151-*bd0739* | Assembly of the following PCR-amplified fragments: opened vector pHCL151 from GL894 with primers oGL1702-1720; *mscarlet-linker* from GL894 with oGL2493-1701; and *bd0739* from HD100 gDNA with primers oGL2531-2532. |
| GL2385: pHCL147-*bd1937* | Assembly of the following PCR-amplified fragments: opened vector pHCL147 from GL891 with primers oGL1696-1720; and *bd1937* from HD100 gDNA with primers oGL2533-2534. |
| GL2386: pHCL147-*bd2402* | Assembly of the following PCR-amplified fragments: opened vector pHCL147 from GL891 with primers oGL1696-1720; and *bd2402* from HD100 gDNA with primers oGL2535-2536. |
| GL2388: pHCL147-*bd3734* | Assembly of the following PCR-amplified fragments: opened vector pHCL147 from GL891 with primers oGL1696-1720; and *bd3734 (mglA)* from HD100 gDNA with primers oGL2539-2540. |
| GL2390: pHCL147-*bd2490* | Assembly of the following PCR-amplified fragments: opened vector pHCL147 from GL891 with primers oGL1696-1720; and *bd2490* from HD100 gDNA with primers oGL2543-2544. |
| GL2391: pHCL147-*bd2492* | Assembly of the following PCR-amplified fragments: opened vector pHCL147 from GL891 with primers oGL1696-1720; and *bd2492 (sgmX)* from HD100 gDNA with primers oGL2545-2546. |
| GL2392: pHCL147-*bd3125* | Assembly of the following PCR-amplified fragments: opened vector pHCL147 from GL891 with primers oGL1696-1720; and *bd3125 (cdgA)* from HD100 gDNA with primers oGL2547-2548. |
| GL2394: pHCL147-*bd3262* | Assembly of the following PCR-amplified fragments: opened vector pHCL147 from GL891 with primers oGL1696-1720; and *bd3262* from HD100 gDNA with primers oGL2551-2552. |
| GL2395: pHCL151-*bd3261* | Assembly of the following PCR-amplified fragments: opened vector pHCL151 from GL894 with primers oGL1702-1720; *mscarlet-linker* from GL894 with oGL2493-1701; and *bd3261* from HD100 gDNA with primers oGL2553-2554. |
| GL2396: pHCL151-*bd3273* | Assembly of the following PCR-amplified fragments: opened vector pHCL151 from GL894 with primers oGL1702-1720; *mscarlet-linker* from GL894 with oGL2493-1701; and *bd3273* from HD100 gDNA with primers oGL2555-2556. |
| GL2397: pHCL147-*bd0164* | Assembly of the following PCR-amplified fragments: opened vector pHCL147 from GL891 with primers oGL1696-1720; and *bd0164 (romY)* from HD100 gDNA with primers oGL2557-2558. |
| GL2398: pTNV215-*msfgfp-bd0739* | Assembly of the following PCR-amplified fragments: opened vector pTNV215 from GL606 miniprep with primers oGL451-1299; *promoter(bd0739)* (without *bd0739*) from HD100 gDNA with primers oGL2171-2146; *msfgfp-4GS* with primers oGL683-350; and *bd0739* from HD100 gDNA with primers oGL2147-2148. |
| GL2399: pTNV215-*bd1937-msfgfp* | Assembly of the following PCR-amplified fragments: opened vector pTNV215 from GL606 miniprep with primers oGL451-1299; *promoter(bd1937)* (including *bd1937*) from HD100 gDNA with primers oGL2174-2173; and *4GS-msfgfp* with primers oGL1316-673. |

### Table S3 – Oligos used in this study

| Primer name | Primer sequence (5’>3’) |
|---|---|
| oGL264 | TCTAGAGTCGACCTGCAG |
| oGL265 | GGATCCCCGGGTACCGAG |
| oGL286 | GGTacccggggatcctcta |
| oGL287 | CGACTCTAGAGGATCCCCGGGTACCcccgggacgctgccgcaa |
| oGL288 | gaattctcctcatcctgtctct |
| oGL299 | GGCTCAGGAAGCGGCTCAGGATCCAAAGGA |
| oGL308 | GGATCCTGAGCCGCTTCCTGAGCCGTTGTTCAAAACCTTGTTGCTCTTGGA |
| oGL322 | GCTAGCCCAAAAAAACGGGTATG |
| oGL331 | ggcactggccgtcgttttacaa |
| oGL332 | gtaatcatgtcatagctgtttcctgtgtgaaa |
| oGL333 | TGAGCCGCTTCCTGAGCCttcagcagaaagaggggacacg |
| oGL334 | cacacaggaaacagctatgacatgattaccggcgatgagaaacaaagaaatgg |
| oGL335 | gtaaaacgacggccagtgccccgccgctttaaagagcaga |
| oGL336 | ACATGGCATGGATGAGCTCTACAAAtaggctctccaaacgtgtcccc |
| oGL337 | ctaTTTGTAGAGCTCATCCATGCCATGTGTAATCC |
| oGL350 | GGATCCTGAGCCGCTTCCTGA |
| oGL399 | ACCCGTTTTTTTGGGCTAGCAGGAGGAATTCACCATGAGCAAAGGAGAAGAACTTTTCA |
| oGL400 | CGCTCTCTGCTGCCCCTGGCTACA |
| oGL451 | ggatcctgatacagattaaatcagaacgcag |
| oGL502 | TCTAGAGTCGACCTGCAGGCA |
| oGL503 | CTCGGTACCCGGGGATCCGGTCGTGGAGTTATCGGT |
| oGL508 | CTGCAGGTCGACTCTAGAAGAAAGCGATTTGAGCTCCGA |
| oGL553 | CAGGAAGCGGCTCAGGATCCAGTACATCAAAACCCAAAGCT |
| oGL554 | GCCTGCAGGTCGACTCTAGACGCCGTCGCCGAGGCTTGGGCGA |
| oGL661 | CGCAGATGACCGCCTTTGCCTGGGA |
| oGL664 | AGTTCTTCTCCTTTGCTCATCCTTAAGTCCTCACGGATGCCTA |
| oGL673 | atttaatctgtatcaggatccctaTTTGTAGAGCTCATCCATGCCAT |
| oGL677 | gcgcggccgcggccta |
| oGL683 | ATGAGCAAAGGAGAAGAACTTTTCA |
| oGL751 | cacacaggaaacagctatgacatgattacgagtacgacagcaaatccat |
| oGL752 | ATCCTGAGCCGCTTCCTGAGCCctgccatccttctttaagcct |
| oGL754 | gtaaaacgacggccagtgccgcattggcggcctgaacag |
| oGL784 | gtcgacctgcaggcatgc |
| oGL897 | GCGGCCGCGTCGTGAC |
| oGL929 | cgcctaggccgcggccgcgcCGGCATTATATGCACCTGTTCTA |

| | |
|---|---|
| **oGL930** | cccaGTCACGACGCGGCCGCctaTTTGTAGAGCTCATCCATGCCA |
| **oGL931** | cgcctaggccgcggccgcgcGTGCGCGTGCGCCCGCACGAAGTGA |
| **oGL932** | cccaGTCACGACGCGGCCGCCGCCGTCGCCGAGGCTTGGGCGA |
| **oGL991** | ATAATacctccttaTctagAGGATCCCCG |
| **oGL992** | AagctTGGCTGTTTTGGCG |
| **oGL993** | TCCTctagAtaaggaggtATTATATGAGAATTACTCCTATCGATATCGCTCAC |
| **oGL994** | CCAAAACAGCCAagctTtaTTTGTAGAGCTCATCCATGCCATGT |
| **oGL1211** | attattcagcagatctcattttgttcctccttgaacaatatga |
| **oGL1212** | gaacaaaatgagatctgctgaataatcagcacgaagaca |
| **oGL1236** | caggaaacagctatgacatgattacAATCCCAAAGCGGAAGACGAA |
| **oGL1238** | ATGCCTGCAGGTCGACTCTACTCAGATTCATGCGGTGATTGACG |
| **oGL1299** | cccgggccagctgcattaat |
| **oGL1316** | GGCTCAGGAAGCGGCTC |
| **oGL1628** | CAGCAGCGGGATCATGGATCCTGAGCCGCTTCCTGAGCCAATGGACTTTTCAGTTTCGCGGA |
| **oGL1629** | ATGATCCCGCTGCTGAACGC |
| **oGL1633** | CAGCAGCGGGATCATGGATCCTGAGCCGCTTCCTGAGCCttcagcagaaagaggggacacg |
| **oGL1696** | tctggtctcgagggtccg |
| **oGL1701** | cagaccagccggagaacc |
| **oGL1702** | GCTTATCGATCTCACGATAATATCCGG |
| **oGL1720** | CATATGTATATCTCCTTCTTAAAGTTAAAC |
| **oGL1729** | TCATTTATCGTCATCGTCTTTGTAGTCCT |
| **oGL1730** | CAAAGACGATGACGATAAATGAATCCTCCGCGAAACTGAAAAGTC |
| **oGL1731** | CAAAGACGATGACGATAAATGAgctctccaaacgtgtcccc |
| **oGL1767** | GGCTCAGGAAGCGGCTCAGGATCCATCCCGCTGCTGAACGC |
| **oGL1768** | TCATTTATCGTCATCGTCTT |
| **oGL1769** | CTACAAAGACGATGACGATAAATGAggtttgagcgatgagcttcagaaaa |
| **oGL2146** | gttcttctcctttgctcatgtctaagggaaatatcggaat |
| **oGL2147** | GAAGCGGCTCAGGATCCatgagcaaaaagaaaaagaaatcttcagccg |
| **oGL2148** | ctgatttaatctgtatcaggatccctcagctttcttcaacggaacgaga |
| **oGL2171** | attaatgcagctggcccgggcgcagggctgtttttgagatgt |
| **oGL2173** | attaatgcagctggcccgggccagatctttcaggtgctgcat |
| **oGL2174** | GAGCCGCTTCCTGAGCCtgccgcctgcagaccttt |
| **oGL2464** | ctcgagggtggaggctc |
| **oGL2465** | catatgtatatctccttCTTAAAtctagacagcg |
| **oGL2485** | agaTTTAAGaaggagatatacatatgGCTTTACGCGTCTTGCTTGC |
| **oGL2486** | gcctccaccctcgagAATGGACTTTTCAGTTTCGCGG |
| **oGL2493** | CTTTAAGAAGGAGATATACATATGatggtttctaaaggtgaagcagttatc |
| **oGL2531** | tctccggctggtctgagcaaaaagaaaaagaaatcttcagccg |

| | |
|---|---|
| **oGL2532** | TTATCGTGAGATCGATAAGCctactcagctttcttcaacggaacga |
| **oGL2533** | TAACTTTAAGAAGGAGATATACATATGAGAACCTCTAAGATAATTTGCCCTTTTAGG |
| **oGL2534** | accctcgagaccagatgccgcctgcagacc |
| **oGL2535** | TAACTTTAAGAAGGAGATATACATATGGAATCAGGCAAATCTAAGATCTTGATATTG |
| **oGL2536** | accctcgagaccagatttctttttaaagaagcccgaaacaacctgtt |
| **oGL2539** | TAACTTTAAGAAGGAGATATACATATGtcctttattaactacaatgccaaagaaattca |
| **oGL2540** | accctcgagaccagaCAGAGTCGTTCCGCCTTTTAGAAC |
| **oGL2543** | TAACTTTAAGAAGGAGATATACATATGCCCAAAATTGAAGCAAGCAC |
| **oGL2544** | accctcgagaccagaACCTTGGGCGCGGTAG |
| **oGL2545** | TAACTTTAAGAAGGAGATATACATATGTCCACATATATTGAGTTAGAAATCCAGA |
| **oGL2546** | accctcgagaccagactggccaccccagatg |
| **oGL2547** | TAACTTTAAGAAGGAGATATACATATGAACATTCGCGATTACAGTTCTCAG |
| **oGL2548** | accctcgagaccagattccgctgtcacttcaaattcagg |
| **oGL2551** | TAACTTTAAGAAGGAGATATACATATGCTTAAAGTCGGACAACTTTTGAAGTTTG |
| **oGL2552** | accctcgagaccagaGTTTATTAAGTCGAACTCTTTCAGAGGAGT |
| **oGL2553** | tctccggctggtctgGAAAAGAAGAATTTTTTAAGTCCTATGGAGGG |
| **oGL2554** | TTATCGTGAGATCGATAAGCTTACAGACGCTTCAGTCGCG |
| **oGL2555** | tctccggctggtctgGATCCTTTTGAGGAATTTGAGTTTAAGC |
| **oGL2556** | TTATCGTGAGATCGATAAGCTTATGCCTTTTTGAACAGCTGAGCC |
| **oGL2557** | TAACTTTAAGAAGGAGATATACATATGCAGGTGCAAAAAGGTTTTAATTCAG |
| **oGL2558** | accctcgagaccagaAGGAAGGCTCCCTGCC |
