## Supplementary figures and images for "Distinct dynamics and proximity networks of hub proteins at the prey-invading cell pole in a predatory bacterium"

### Supp Figures 1 to 5

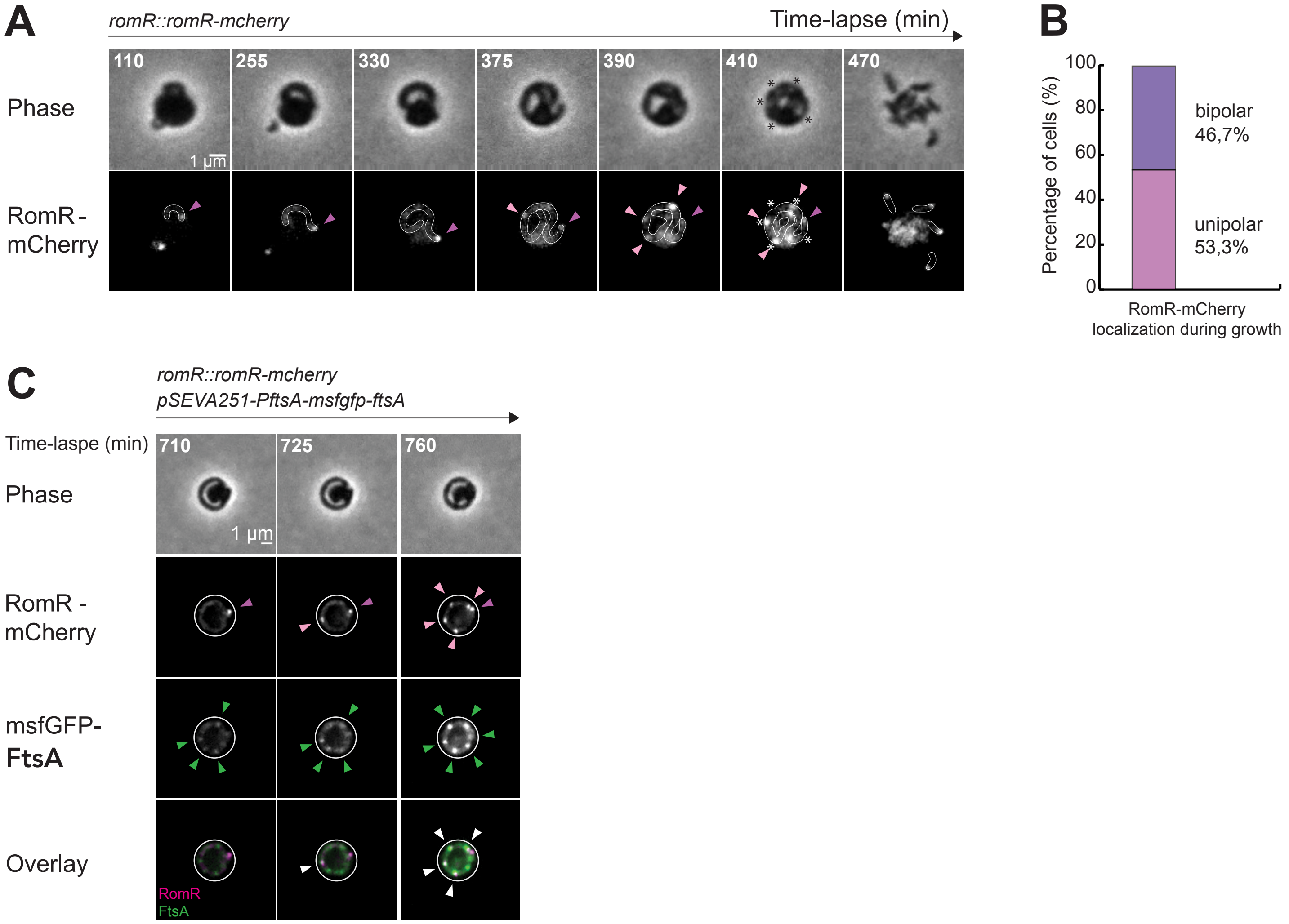

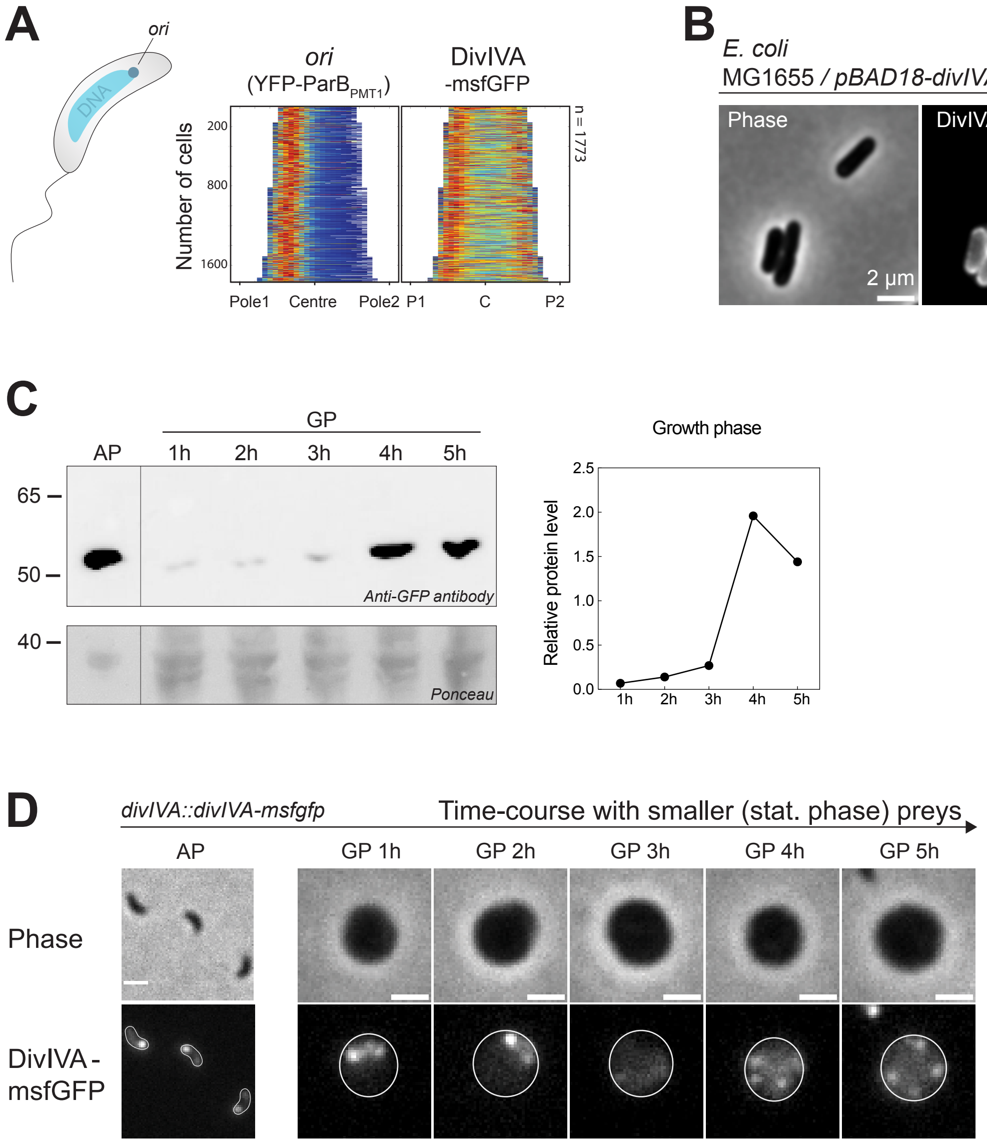

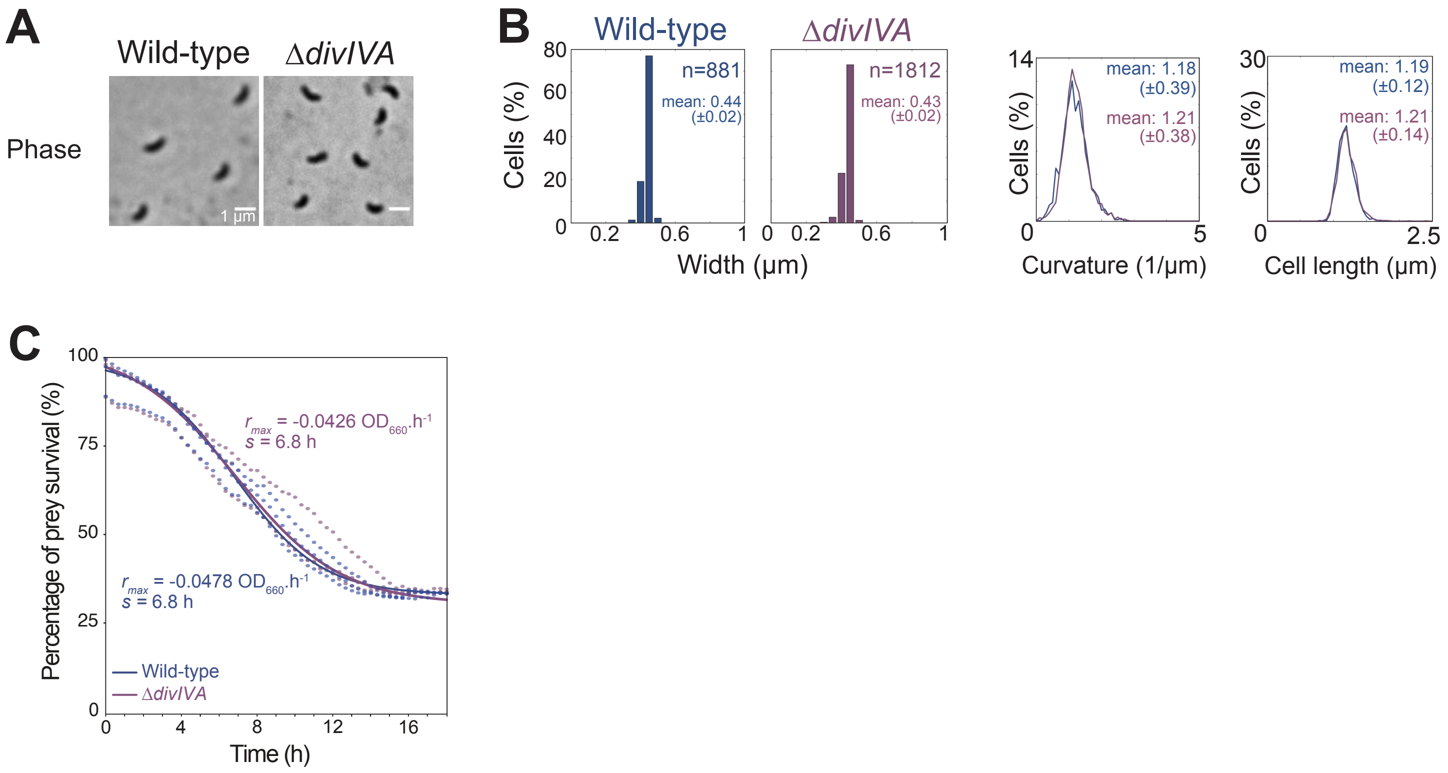

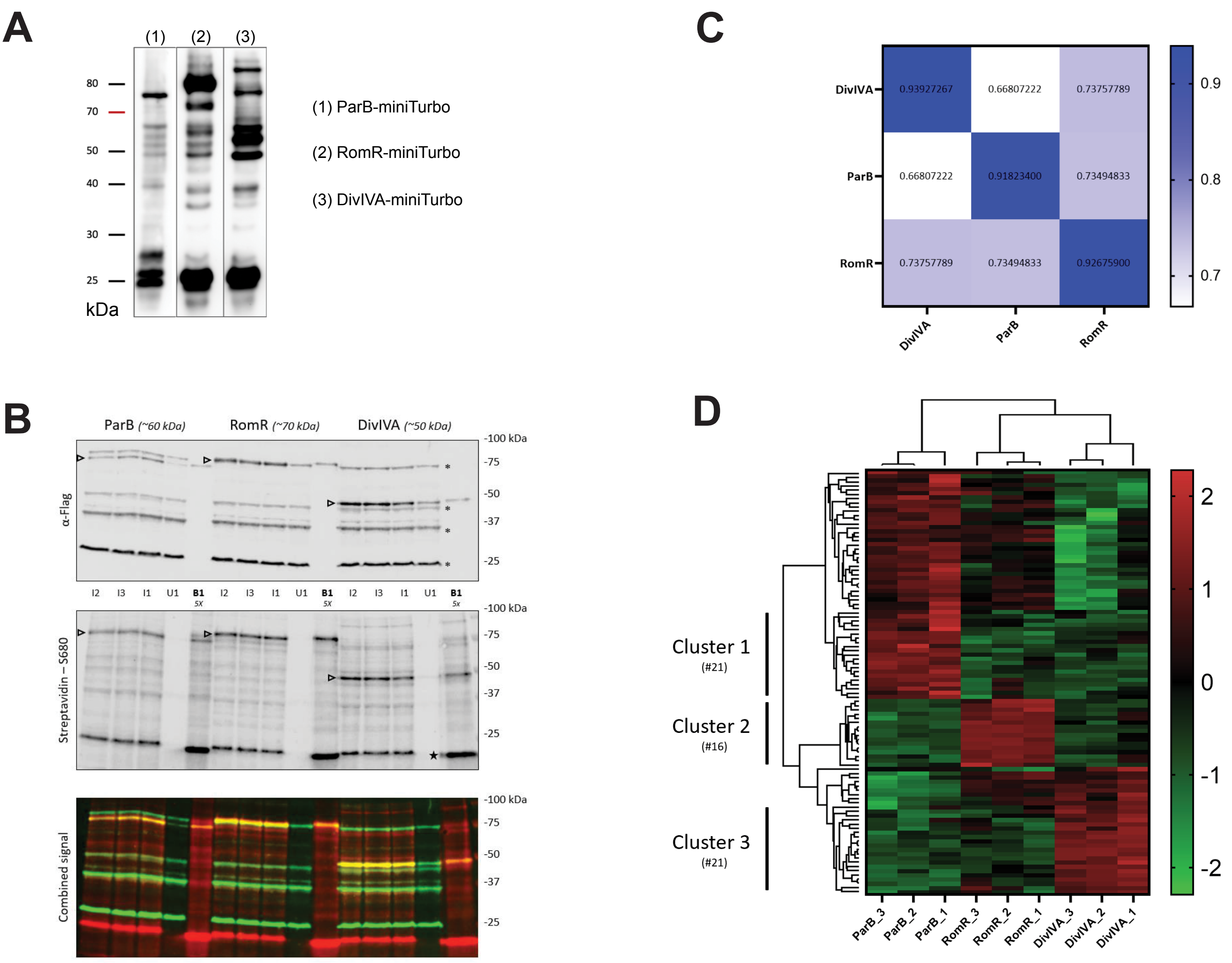

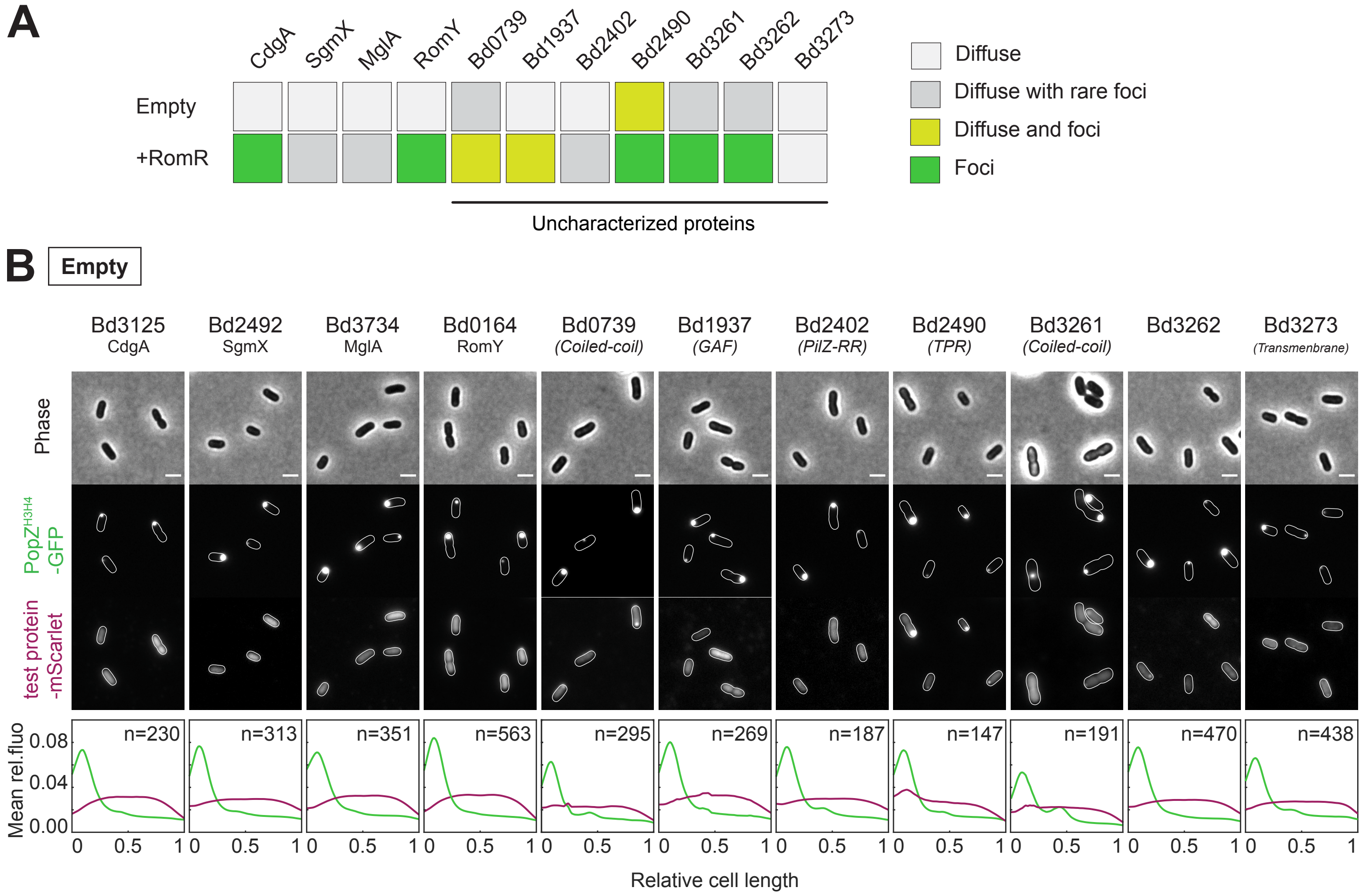
